# On correctness of gene tree tagging under a unified model of gene duplication, loss, and coalescence

**DOI:** 10.64898/2026.01.20.700722

**Authors:** Rachel A. Parsons, Yunzhuo Liu, Parth Dua, Alexey Markin, Erin K. Molloy

## Abstract

ASTRAL-pro is the leading method for reconstructing species trees under complex evolutionary scenarios involving gene duplication, loss, and coalescence, commonly modeled by DLCoal. A unique aspect of A-pro is that it utilizes rooted gene trees, with internal vertices labeled as duplications or speciations, to modify its objective function compared to the traditional ASTRAL method. Although there is a natural event-based definition of correct tagging when genes evolve with only gene duplications and losses, it cannot be applied when there is deep coalescence. Here, we introduce a definition of correct tagging that is broadly applicable, proposing that a gene tree vertex is correctly tagged as a duplication if it is the most recent common ancestor of at least one pair of gene copies related via a duplication event. Using this definition, we study some statistical properties of ASTRAL-pro’s objective function under the DLCoal model and evaluate the accuracy of ASTRAL-pro’s tagging algorithm in simulations.

## Introduction

The availability of genome-scale data sets has increased dramatically over the last decade, thanks to technological advances and large-scale initiatives, like One Thousand Plant Transcriptomes [Leebens-Mack et al., 2019] and Bird 10K [Feng et al., 2020]. Fully leveraging these data for species tree reconstruction requires accounting for gene tree heterogeneity (GTH), which broadly refers to variation in the evolutionary histories across different genomic regions, called genes [Maddison, 1997]. Perhaps the most pervasive source of GTH is incomplete lineage sorting (ILS) [Edwards, 2009], where gene trees disagree with the species tree due to deep coalescence, a population-genetics phenomenon, modeled by the Multi-Species Coalescent (MSC) [Pamilo and Nei, 1988, Rannala and Yang, 2003]. Under the MSC, the most probable gene tree agrees with the species tree for unrooted four-taxon trees, called quartets [Allman et al., 2011]. This result motivates the most popular species tree estimation method, ASTRAL, which seeks a species tree that maximizes the number of quartets also induced by the input gene trees, estimated from molecular sequences [Mirarab et al., 2014].

ASTRAL’s popularity may be attributed to its practical properties (speed and accuracy) as well as its statistical guarantee; specifically, ASTRAL is *consistent* under the MSC, meaning that it returns the true unrooted species tree topology with high probability, as the number of gene trees generated under the MSC tends to infinity. An underlying assumption is that genes evolve without duplications. Thus, multi-copy gene families must be restricted to orthologs or else excluded from species tree reconstruction. The former can present computational challenges (see Majidian et al., 2025 for a recent review), and the latter can dramatically reduce the amount of data available for species tree reconstruction (consider that Wickett et al., 2014 curated more than 9,000 gene families but just 424 were single-copy). To address this issue, some methods reconstruct species trees directly from multi-copy gene trees. However, the accuracy of traditional approaches based on gene tree parsimony (e.g., Parsons and Bansal, 2025) or Robinson-Foulds supertrees (e.g., Molloy and Warnow, 2020) is challenged by deep coalescence as well as gene tree estimation error (Fig. 2b in Zhang et al., 2020).

**Fig. 1.**
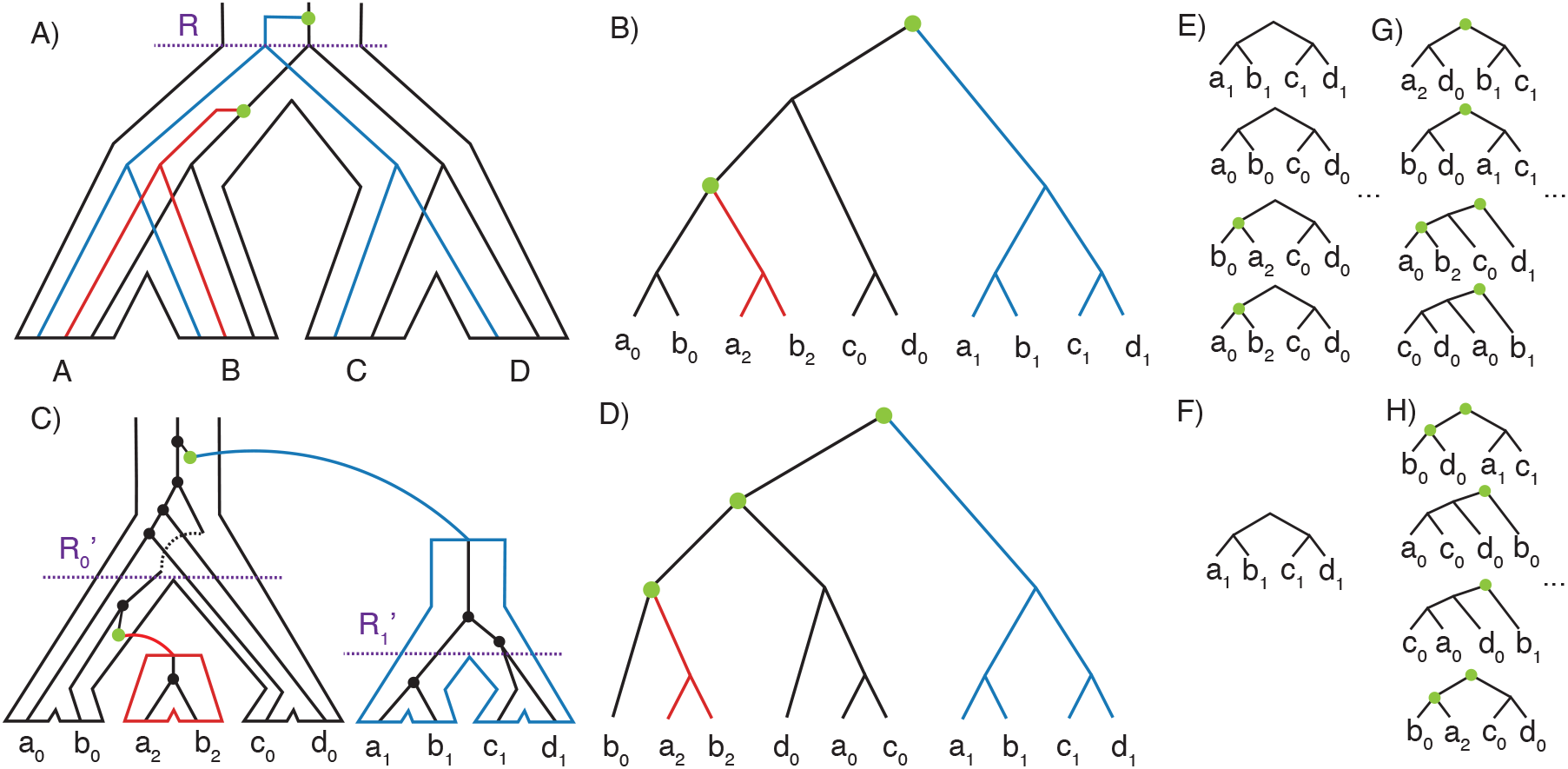
**A)** Locus tree evolving within species tree, with duplications (no losses shown). **B)** Correctly tagged locus tree with green circles indicating duplication events. **C)** Gene tree evolving within locus tree, with deep coalescence (branch lengths/widths of locus tree are not drawn to scale). Black circles represent coalescent events. **D)** Correctly tagged gene tree with green circles indicating tagged duplication vertices. **E)** There are 3 · 3 · 2 · 2 = 36 ways to select four gene copies (i.e., leaves) uniquely labeled by *A, B, C, D*. Five induce SQs in the locus tree, corresponding to the ways of selecting four gene copies that descend from the same locus lineage at *R*, shown in black and blue. The SQ on *a*_2_, *b*_2_, *c*_0_, *c*_0_ is <u>not</u> shown. **F)** Because of deep coalescence, just one of the five SQs in the locus tree is also an SQ in the gene tree. **G)** and **H)** Example DQs induced by the locus tree and gene tree, respectively.

**Fig. 2.**
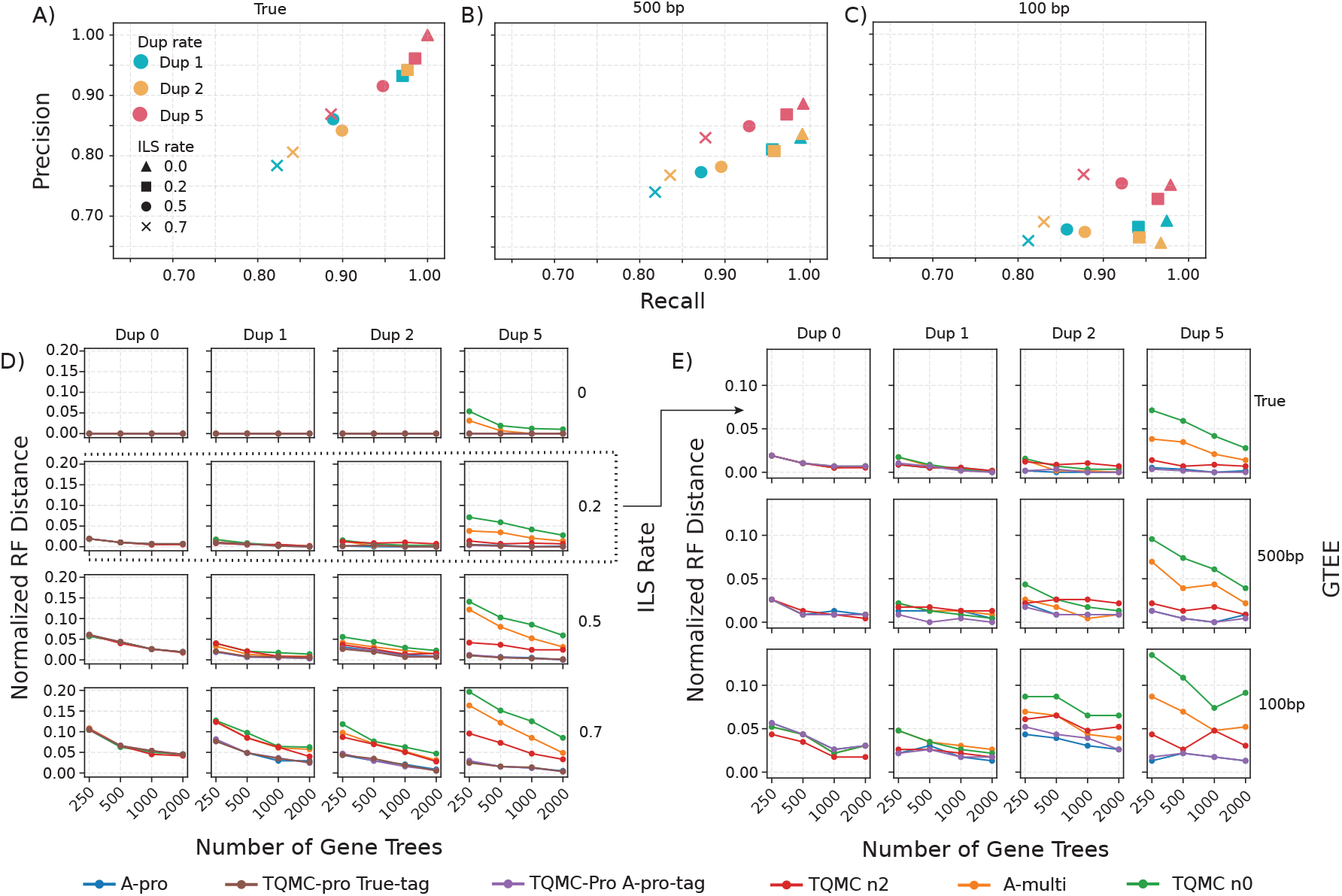
**A, B, and C** Precision and recall of A-Pro’s tagging algorithm on true gene trees and gene trees estimated 500bp and 100bp, respectively. **D)** Species tree error varying duplication and ILS levels (loss rate=0). **E)** Species tree error varying GTEE (ILS level=0.2). Dots are averages over 25 replicates for true gene trees and 10 replicates for estimated gene trees.

The current leading methods are based on quartets. ASTRAL-multi (called A-multi for brevity) extends ASTRAL’s objective function to multi-copy genes in the natural way [Rabiee et al., 2019]. Although A-multi is statistically consistent under a unified duplication-loss-coalescent model, called DLCoal [Markin and Eulenstein, 2021, Hill et al., 2022], it is less accurate than a related method, called A-pro, in simulations (Figs. 2b and 4 in Zhang et al. [2020]), for which there is currently no proof of consistency. Unlike A-multi, A-pro internally roots and *tags* gene trees so that each internal vertex is labeled as duplication or speciation. This information is used to *exclude* the score contributions from quartets induced by duplication vertices, called “duplication quartets”, and to *agglomerate* or combine the score contributions from “homoemorphic speciation quartets” quartets into a single unit. Tagging these events is natural when genes evolve with gene duplication and loss (GDL) only but not with deep coalescence, as previously noted by Zhang et al. [2020].

Here, we propose a definition of correct tagging that is broadly applicable, proposing that a gene tree vertex is correctly tagged as a duplication if it is the most recent common ancestor of *at least one* pair of gene copies related via a duplication event. Using this definition, we study some statistical properties of A-pro’s objective function under the DLCoal model and evaluate the accuracy of A-pro’s tagging algorithm in simulations.

## Background

We begin with a review terminology and background for phylogenetic methods and the DLCoal model.

### Terminology

A *phylogenetic tree T* is a tuple (g, 𝒳, *ϕ*), where g is a tree in the graph-theoretic sense, 𝒳 is a set of species labels (also called taxa), and *ϕ* is a map from leaves of g to species labels in 𝒳. We say that *T* is *singly-labeled* if *ϕ* is a bijection; otherwise, we say that *T* is *multi-labeled*. For simplicity, we use lower case letters to denote the leaves of gene and locus trees and upper case letters to denote their respective species labels (i.e., *ϕ*(*a*) = *A*). Sometimes we use subscripts (e.g., *a*_0_, *a*_1_, …) to indicate different leaves with the same species label. We let *L*(*T*) denote the leaf set of *T* .

A phylogenetic tree *T* is *rooted* if there exists a directed path from a special vertex with in-degree zero, called the root, to each leaf; it is *unrooted* if all edges are undirected. Non-terminal edges are called internal edges, and non-leaf vertices are called internal vertices. Trees are assumed to be *binary*, meaning that every internal vertex except the root has degree 3 (note that the root vertex has degree 2, otherwise degree 2 vertices are prohibited).

A **quartet** is an unrooted phylogenetic tree with four uniquely labeled leaves. There are three possible quartet topologies indicated by the one internal edge that splits the species set in half; for example, *q*_1_ = *A, B*|*C, D, q*_2_ = *A, C*|*B, D*, and *q*_3_ = *A, D*|*B, C* are the three quartet topologies on species set {*A, B, C, D*}. A quartet can be rooted at its internal branch, which yields a **balanced tree**, or at one of its four terminal branches, which yields **pectinate tree** (sometimes called a caterpillar tree). A larger tree *T induces* a collection of quartets that can be computed by restricting *T* to every subset of four uniquely labeled leaves (i.e., deleting all other leaves and suppressing vertices of degree 2). Singly-labeled trees can be restricted according to their species labels as the outcome is unambiguous. If *T* is rooted, the induced subtree on *A, B, C, D*, denoted *T* |_*A,B,C,D*_, inherits the rooting, although it is ignored when convenient and clear from context.

### DLCoal Model

The DLCoal model, introduced by Rasmussen and Kellis [2012], is parameterized by a **species tree** *T* on taxon set 𝒳. Species trees must be rooted, binary, singly-labeled. Each branch *e* of *T* is annotated with length *t*(*e*), indicating the number of generations, as well as width *N*_*e*_(*e*), indicating the effective population size. Additionally, duplication and loss rates, *λ* and *µ* respectively, must be specified (rates correspond to the number of events per generation per lineage). Gene trees evolve within a species tree under duplication, loss, and coalescent processes in two steps, which we summarize next (see Hill et al., 2022 for details).

First, a **locus tree** tree *ℓ* evolves within the species tree *T* according to the stochastic GDL model of Arvestad et al. [2009] (Fig. 1A–B). A single lineage starts at the root of *T* and grows forward in time. Lineages are duplicated and lost on the branches of *T* with rate *λ* and *µ*, respectively. A **duplication event** instantaneously splits the impacted lineage, with one side being arbitrarily designated as the original (i.e., mother) copy and the other being designated as the duplicated (i.e., daughter) copy. A loss event prohibits the impacted lineage from growing forward (it is later pruned from *ℓ*). When a lineage arrives at an internal node *v* of *T*, it undergoes a **speciation event**, instantaneously splitting so that one side enters the branch *v* ↦ *v*.*l* and the other side enters the branch *v* ↦ *v*.*r*, where *v*.*l* and *v*.*r* indicate the left and right child of *v*, respectively. Once a lineage arrives at a leaf vertex of *T*, it inherits the associated taxon label; likewise, branches in *ℓ* inherit lengths and widths from *T* .

Next, a **gene tree** *g* evolves within the locus tree *ℓ* under the Multilocus Coalesecent (MLC) (Fig. 1C–D), which is closely related to the MSC. Lineages sampled at leaves of *ℓ* trace their ancestry backward in time. Multiple lineages entering same branch of *ℓ* can coalesce with equal probability; the resulting internal vertices represent **coalescent events**. The *bounded coalescent* is used to force gene copies to coalesce prior to the duplication event from which they originate (e.g., gene copies *a*_2_ and *b*_2_ in Fig. 1C–D). Lineages entering the branch above the root of *ℓ* will eventually coalesce into a single lineage [Rannala and Yang, 2003].

To summarize, the DLCoal model, given species tree *T* on taxon set 𝒳, generates a gene-locus tree pair, denoted *ℓ, g*∼*DLCoal*(*T*, 𝒳), although only *g* is observed. Because of this generation process, *ℓ, g* are rooted and binary and can be multi-labeled. One lineage is sampled per leaf in *ℓ*, so there is a bijection between the leaf sets of *ℓ* and *g* (so we reference both of them with the variables, typically *a, b, c, d*, which are labeled *A, B, C, D*, respectively). Any collection of gene trees generated under DLCoal is assumed to be independent and identically distributed.

### Maximum Quartet Support Species Trees (MQSST)

ASTRAL is a heuristic for the NP-hard MQSST problem [Lafond and Scornavacca, 2019], defined by the following input/output:

- **Input**: a set 𝒯 of *k* unrooted, binary trees, each with leaves labeled by species set *X*
- **Output**: an unrooted, binary tree with leaves bijectively labeled by *X* that maximizes the number of quartets also displayed by the input gene trees

Likewise, A-multi is a heuristic for MQSST but extended to multi-labeled input trees in the natural way. Let *N*^*t*^(*T* |_*A,B,C,D*_) be <u>number</u> of quartets in a multi-labeled input tree *t* that agree with some singly-labeled tree *T* restricted to *A, B, C, D*, that is

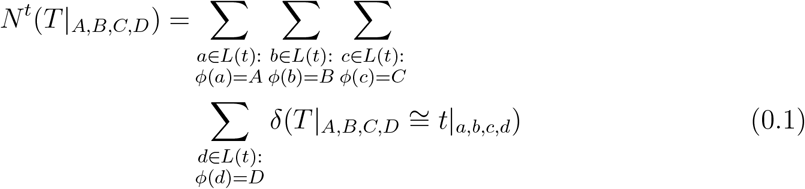

where *δ*(·) returns 1 if the two quartets are isomorphic and 0 otherwise. Then, A-multi’s objective function is

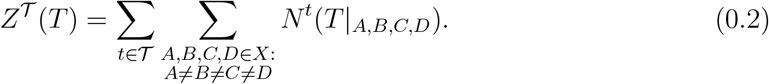

It is easy to see that A-multi’s objective function is equivalent to ASTRAL’s when the input trees are singly-labeled.

As previously mentioned, the leading method A-pro is similar to A-multi but (1) *excludes* “duplication quartets” and (2) *agglomerates* “speciation quartets” that are homeomorphic up to duplications. A quartet is classified as a **speciation quartet** (SQ) if for every selection of three leaves, their most recent common ancestor (MRCA) is a speciation vertex (Fig. 1E–F); otherwise, the quartet is classified as a **duplication quartet** (DQ) (Fig. 1G–H). Thus, a balanced quartet is an SQ if the root (i.e., the MRCA of its leaves) is a speciation vertex; in contrast, a pectinate quartet is an SQ if both the root and the MRCA of its three non-outgroup leaves are speciation vertices. To classify quartets as SQs or DQs, the input trees must be rooted and **tagged** so that their internal vertices are labeled as duplications or speciations. For convenience, A-pro implements its own tagging algorithm and roots each input tree such that the GDL score associated with tagging is minimized.

The motivation behind A-pro is that (1) quartets whose topologies are driven by duplication vertices do not represent information about speciation events and thus should be excluded and (2) isomorphic quartets whose topologies are driven by the same speciation vertices (but whose leaves differ due to duplications) do not provide new information and thus should be counted as a single unit [Zhang et al., 2020]. Henceforth, we focus on a version of A-pro’s objective function that counts SQs only, denoted by subscript *sq* (e.g., *δ*_*sq*_, *N*_*sq*_, and *Z*_*sq*_); we leave agglomeration of homeomorphic SQs to future work.

## Methods

### Correctness of Gene Tree Tagging under DLCoal

As previously mentioned, when genes evolve with GDL only, there is a natural definition of correct tagging. Take Figure 1A–B, in which locus tree *ℓ* evolves within species tree *T*, as an example. Consider the internal vertex *v* in locus tree *ℓ* associated with the duplication event at the MRCA of *A, B* in *T* . The leaves below *v*.*l* (i.e., *a*_0_, *b*_0_) are descendants of the original (i.e., mother) copy, and those below *v*.*r* (i.e., *a*_2_, *b*_2_) are descendants of the duplicated (i.e., daughter) copy. Any pair of leaves with MRCA *v* descend from the same duplication event and thus are **paralogs**.

This argument breaks down with deep coalesence. Take Figure 1C–D, in which gene tree *g* evolves within locus tree *ℓ*, as an example. Consider the internal vertex *u* in gene tree *g* with leaves *b*_0_, *a*_2_, *b*_2_ descending from *u*.*l* and leaves *a*_0_, *c*_0_, *d*_0_ descending from *u*.*r*. Looking at the collection of leaf pairs with MRCA *u*, some are related via a duplication event (e.g., *a*_2_ and *a*_0_) but others are not (e.g., *b*_0_ and *a*_0_). The latter occurs because of deep coalescence (e.g., both *a*_0_ and *c*_0_ fail to coalesce until they reach a deeper ancestral population, after which they coalesce with each other). To summarize, whether or not *u* should be tagged as a duplication vertex depends on the gene copies being examined; however, a single tag must be designated. We propose the following conservative approach.

Definition 1 (Correct Duplication Tagging for Gene Trees) Let *ℓ, g* be a locus-gene tree pair, and let *u* be the internal node in *g*. We say that *u* is correctly tagged as a **duplication** if there exists one leaf *x* below *u*.*l* and one leaf *y* below *u*.*r* such that *x, y* are paralogs (i.e., the MRCA of *x, y* in *ℓ* is a duplication vertex).

Our definition of correct tagging has two advantages. First, it is backwards compatible with locus trees (i.e., applying our definition of correct tagging to a locus tree successfully captures the duplication events). Second, it is well-aligned with A-pro’s tagging algorithm, which would designate *u* as a duplication vertex because it is the MRCA of two gene copies *a*_0_ and *a*_1_ in the same species *A* so they must be related via a duplication event. It is worth noting that A-pro’s algorithm can fail to correctly tag gene trees (and locus trees) under our definition. As an example, there can be adversarial patterns of duplications and losses that produce single-copy locus subtrees, referred to as adversarial GDL by Molloy and Warnow [2020] (also see Fig. 2D in Molloy and Warnow, 2020). Adversarial GDL would mislead A-pro’s tagging algorithm; similar issues could also arise due to deep coalescence.

It is worth noting that under our definition quartets on **orthologs** can be correctly tagged as DQs because of deep coalescence (e.g., *a*_0_, *b*_0_, *c*_0_, *d*_0_ in Fig. 1H). Conversely, SQs are not restricted to orthologs, even for locus trees (e.g., the quartet on paralogs *a*_2_, *b*_0_, *c*_0_, *d*_0_ is an SQ in Fig. 1E). However, the authors of A-pro highlighted that DQs and SQs need not be equivalent to paralogs or orthologs; therefore, we do not view this as a limitation of our definition of correctness [Zhang et al., 2020].

### Conjecture of A-pro’s statistical consistency under DLCoal and correct tagging

We conjecture that A-pro is statistically consistent under the DLCoal model with the assumption of correct tagging, and provide partial proofs of this conjecture. This conjecture is based on previous simulation results of Zhang et al., 2018.

#### Conjecture 1

An optimal solution under A-pro’s objective function is a consistent estimator of the (unrooted) species tree (topology) under the DLCoal model, assuming the input gene trees are correctly tagged (we implicitly assume no estimation errors or errors in rooting).

We considered whether progress could be made in proving this conjecture based on our definition of correctness, yielding some preliminary results and identifying some key hurdles, which we now describe.

To prove the conjecture, we need to show that the optimal solution under A-pro’s objective function *T* ^∗^ is isomorphic to *T* with high probability, as |𝒫| tends to ∞, where 𝒫 represents a set of gene trees that are independent and identically distributed under the DLCoal model. Following the structure of the proof of Theorem 3 in Legried et al. [2021], let 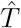 be an arbitrary species tree on 𝒳. By gene tree independence and the law of large numbers, 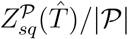 converges almost surely to its expectation simultaneously for all species trees on *X*. Thus, its expectation can be simplified as

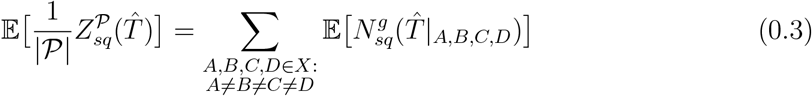

where *g* is an arbitrary tree in 𝒫, that is *g* ∼ *DLCoal*(*T*, 𝒳), and 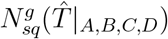 is a random variable for the <u>number</u> of SQs induced by gene tree *g* that are isomorphic to 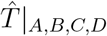 (Equation 0.1). Then, 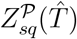 will be maximized by the true species tree *T* if for an arbitrary subset of four species *A, B, C, D* in 𝒳, the following inequality holds

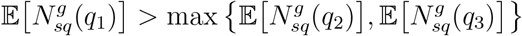

where *T* |_*A,B,C,D*_ = *q*_1_ = *A, B*|*C, D*, without loss of generality, and *q*_2_, *q*_3_ are the alternative topologies. As we will show, there are obstacles to proving this inequality holds.

First, consider how gene copies descend from different lineages in the locus tree *ℓ*, called **locus lineages**, at different vertices of the species tree *T*, similar to Markin and Eulenstein [2021] and Hill et al. [2022]. Let *l*_*X*_ denote the set of locus lineages that exist at vertex *X* in *T*, and let 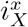 denote the locus lineage in *l*_*X*_ from which gene copy *x* descends. If there are multiple locus lineages in *l*_*X*_, they must descend from duplication events above *X*, so going backward in time, they join at duplication events, eventually becoming one lineage.

Take Figure 1A, in which a locus tree *ℓ* evolves within a balanced species tree *T*, as an example. Let *R* denote the MRCA of *A, B, C, D* in *T* . There are two locus lineages at *R* colored black and blue. Gene copies *a*_0_, *a*_2_, *b*_0_, *b*_2_, *c*_0_, *d*_0_, which descend from the black locus lineage, correspond to the original locus, whereas gene copies *a*_1_, *b*_1_, *c*_1_, *d*_1_, which descend from the blue locus lineage, correspond to the duplicated locus. For any four gene copies *a, b, c, d*, we can consider the relationship among their locus lineages at *R*. As an example, the locus lineage relationship at *R* for *a*_0_, *b*_2_, *c*_1_, *d*_1_ is 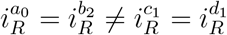, also denoted *ab, cd*. Locus lineage relationships, also called root lineage scenarios, are enumerated by Markin and Eulenstein [2021] and Hill et al. [2022] and then used to partition the computation of 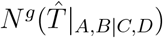 (Equation 0.1). As we will show, locus lineage relationships are also useful for classifying gene tree induced quartets as SQs and DQs.

We now analyze locus lineage relationships, moving down the species tree *T* . To begin, consider locus lineages from **duplication events above the MRCA of** *A, B, C, D*, **denoted** *R*.

##### Lemma 0.1

Assume *ℓ, g* ∼ *DLCoal*(*T*, 𝒳) are correctly tagged. For an arbitrary subset of four taxa {*A, B, C, D*} ⊆ 𝒳, if gene copies *a, b, c, d* do <u>not</u> descend from the same locus lineage at *R*, then *ℓ*|_*a,b,c,d*_ and *g*|_*a,b,c,d*_ are DQs.

*Proof*. Let 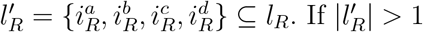, the locus lineages in 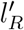 must descend from duplication events above *R*. Therefore, the MRCA of *a, b, c, d* in *ℓ* is a duplication event above *R*, and *ℓ*|_*a,b,c,d*_ is a DQ (this is closely related to the proof of A-pro’s consistency under the GDL-only model; Zhang et al. [2020]). Now consider the MRCA of *a, b, c, d* in *g*, denoted *u*. If 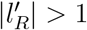, it is always possible to select one gene copy *x* ∈ *L*(*u*.*l*) ∩ {*a, b, c, d*} and one gene copy *y* ∈ *L*(*u*.*r*) ∩ {*a, b, c, d*} such that *x, y* descend from different locus lineages in 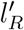 Then, by our definition of correct tagging, *u* is a duplication vertex; it follows that *g*|_*a,b,c,d*_ is a DQ.

If the species tree *T* is pectinate, assume topology (((*A, B*), *C*), *D*) without loss of generality, we must also consider locus lineages from **duplication events below** *R* **but above the MRCA of** *A, B, C*, **denoted** *P* .

##### Lemma 0.2

Assume *T* |_*A,B,C,D*_ = (((*A, B*), *C*), *D*). If gene copies *a, b, c, d* descend from the same locus lineage at *R* but *a, b, c* do <u>not</u> descend from the same locus lineage at *P*, then *ℓ*|_*a,b,c,d*_ and *g*|_*a,b,c,d*_ are DQs.

*Proof*. Suppose gene copies *a, b, c, d* descend from the same locus lineage *r* at *R*. Locus lineage *r* instantaneously splits at *R* in a speciation event that (1) corresponds to the MRCA of *a, b, c, d* in *ℓ* and (2) forces *ℓ*|_*a,b,c,d*_ to be pectinate with outgroup *d*. For *ℓ*|_*a,b,c,d*_ to be a DQ, the MRCA of *a, b, c* must be a duplication vertex. Let 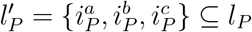. If 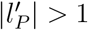, the locus lineages in 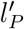 must descend from duplication events above *P* but below *R*, as they must join together prior to the speciation event at *R*. This gives us our result. Due to deep coalescence, we cannot make assumptions about the rooted topology of *g*|_*a,b,c,d*_. There are three cases to consider (1) balanced, (2) pectinate with an outgroup from *a, b, c*, or (3) pectinate with outgroup *d*. For cases (1) and (2), consider the MRCA of *a, b, c, d* in *g*, denoted *u*. If 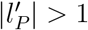, it is always possible to select one gene copy *x* ∈ *L*(*u*.*l*) ∩ {*a, b, c*} and one gene copy *y* ∈ *L*(*u*.*r*) ∩ {*a, b, c*} such that *x, y* descend from different locus lineages in 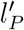. Then, by our definition of correct tagging, *u* is a duplication vertex; it follows that *g*|_*a,b,c,d*_ is a DQ for cases (1) and (2). The same logic holds in case (3) but setting *u* to be the MRCA of *a, b, c* in *g*.

By Lemma 0.1, only gene copies that descend from the same locus lineage at *R* or at *P* can be SQs for the **balanced species tree ((A**,**B)**,**(C**,**D))**. Thus, to prove conjecture 1, it suffices to show

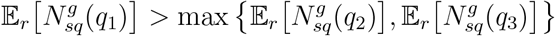

where subscript *r* denotes conditioning the probability space and the expected values on having gene copies from *A, B, C, D* descending from the same locus lineage *r* at *R* (e.g., Markin and Eulenstein [2021], Hill et al. [2022]). Because locus lineage *r* instantaneously splits at *R* in a speciation event, which is the MRCA of *a, b, c, d* in *ℓ*, denoted *R*^′^, we can also condition based on one of two events: (1) no coalescence among *a, b, c, d* below *R*^′^, denoted 𝒩𝒞, and (2) its complement 𝒩𝒞^*C*^ (see Hill et al. [2022]).

#### Coalescence below R^′^

For event 𝒩𝒞^*C*^, either *a, b* coalesce below *R*^′^, *c, d* coalesce below *R*^′^, or both. Regardless, *g*|_*a,b,c,d*_ is isomorphic to *q*_1_ = *A, B*|*C, D*, so 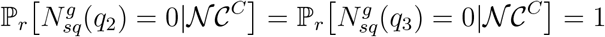. Although the unrooted topology of *g*|_*a,b,c,d*_ is fully determined by our conditioning, it could be either an SQ or a DQ, depending on *g* (because complex scenarios involving duplications and deep coalescence, even among gene copies outside *a, b, c, d*, can trigger duplication vertices across *g*; see Fig. 1). However, we know that *ℓ*|_*a,b,c,d*_ must be an SQ with rooted topology ((*a, b*), (*c, d*)) by our assumption that all gene copies descend from *r*, and we know that there is a strictly positive probability of *g* evolves within *ℓ* under the MLC without deep coalescent events, in which case *g* is isomorphic to *ℓ*. This gives us 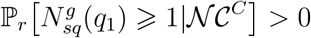, and by Markov’s Inequality, 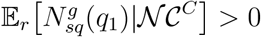, which gives

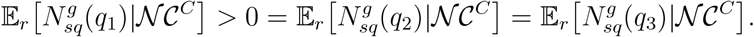

At this point, we should need to consider the issue of homeomorphic SQs being counted as a single unit. To simplify our discussion, we ignore this aspect of A-pro’s objective function, effectively using a version that excludes DQs only (and counts all SQs).

#### No coalescence below R^′^

For event 𝒩𝒞, gene copies *a, b, c, d* fail to coalesce and enter above *R*^′^, so all three quartet topologies are possible. To prove the conjecture for the balanced species tree under an exclusion only version of A-pro’s objective function, we need to show that 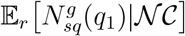 is at least large as 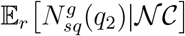 and 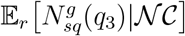. This inequality is traditionally argued based on the exchangeability of lineages after they enter a shared ancestral population; however, in our case, we can not necessarily treat lineages as exchangeable as lineage swapping can trigger/untrigger duplication vertices, ultimately changing whether the induced quartet on *a, b, c, d* is an SQ or a DQ, even for our definition of correct tagging, which up until now has been convenient.

Consider an adversarial scenario where locus tree *ℓ* evolves within species tree ((*A, B*), (*C, D*)) with no duplications in the ancestor of *A, B*, (2) no duplications in the ancestor of *C, D*, and (3) duplications in one or more of the terminal branches of *A, B, C, D* with the duplication events always acting on the mother locus. Take locus tree *ℓ* = ((*a*_0_, *b*_0_), ((*c*_0_, *c*_1_), *d*)) and gene tree *g* = ((*c*_0_, (*a*_0_, *c*_1_)), (*b*_0_, *d*_0_)) as an example of this scenario (and note that *a*_0_, *b*_0_ and *c*_0_, *d*_0_ could not have coalesced below *R*^′^). The induced quartet is an SQ that disagrees with the species tree (*a*_0_, *c*_0_|*b*_0_, *d*_0_). Lineages *b*_0_ and *c*_0_ are exchangeable under the coalescent in this example; however, swapping them yields tree ((*b*_0_, (*a*_0_, *c*_1_)), (*c*_0_, *d*_0_)), which occurs under the DLCoal model with the same probability as *g*, but now the induced quartet is a DQ that agrees with the species tree (*a*_0_, *b*_0_|*c*_0_, *d*_0_). More broadly, under this adversarial scenario,

1. all gene copies can enter above *R*^′^ (because duplications acted only on the mother locus so they are not governed by the bounded coalescent);
2. any selection of *a, b, c, d* may form an SQ in *g* (because duplications occurred only on the terminal branches so any selection of *a, b, c, d* are orthologs and are not related through a duplication vertex in *ℓ*);
3. swapping lineages *a, b, c, d* has the potential to trigger/untrigger duplication (based on the particular arrangement of *a*’s in *g* and similarly for *b, c, d*).

Therefore, a more delicate argument is needed to prove the inequality, so the consistency of A-pro is an open question.

A similar scenario occurs for the pectinate tree **(((A**,**B)**,**C)**,**D)**, except that only gene copies that descend from the same locus lineage at *R* and *P* can be SQs, by Lemmas 0.1 and 0.2. Thus, we would instead condition the probability space and expected values on having gene copies from *A, B, C, D* descending from the same locus lineage *r* at *R* with those from *A, B, C* descending from the same locus lineage *p* at *P* .

Lastly, it is worth noting that we did not consider the duplication rate *λ* or loss rate *µ*, instead treating the GDL model generically (e.g., Molloy and Warnow [2020]). One possibility is that specific settings of these and other numerical parameters (i.e., species tree branch lengths and widths) could make adversarial scenarios, like the one we described above, have a low probability under the DLCoal model. We leave further exploration of this topic to future work.

### TREE-QMC-pro

Next, we empirically evaluate the exclusion-only version of A-pro’s objective function by implementing it within TREE-QMC [Han and Molloy, 2023]. TREE-QMC (or TQMC for brevity) is a heuristic for the same NP-hard optimization problem as ASTRAL but is instead based on the Quartet Max Quartet (QMC) framework of Snir and Rao [2010]. QMC reconstructs the species tree in a divide-and-conquer fashion. At each step in the divide phase, the goal is to partition the taxon set in half, producing an internal branch of the output species tree. Each (bi)partition is identified by constructing a graph in which vertices are the taxon set for the given subproblem and edges are weighted according to the input quartets. Each quartet *A, B*|*C, D* contributes two bad edges (i.e., (*A, B*) and (*C, D*)) and four “good” edges (i.e., (*A, C*), (*A, D*), (*B, C*), and (*B, D*)). Cutting the bad edges is undesirable because it breaks apart siblings, whereas cutting the good edges is desirable; this motivates seeking a maximum cut; see Snir and Rao [2010].

The main contribution of TREE-QMC is that it constructs the quartet graph in an efficient manner directly from gene trees, without enumerating all quartets. The latest version enables quartets to be weighted based on gene tree branch lengths and support values to improve robustness to gene tree estimation error [Han and Molloy, 2025]. To build the weighted graph efficiently, TREE-QMC uses “auxiliary values” that store information about how taxa below any given node can be combined to produce good or bad edges. Auxiliary values are computed in either preorder or postorder traversals of the input gene tree; after, good/bad edges are computed in postorder traversals. Our contribution is modifying the recurrences for auxiliary values and good/bad edges to exclude contributions from DQs (see Section 1 of the Supplement for details and equations). To our knowledge, our new method, called TQMC-pro, is the first to enable both the exclusion of DQs and the weighting of quartets based on gene tree branch information. Including the agglomeration in this framework is challenging because homeomorphic SQs can have different weights.

Lastly, the other contribution of TREE-QMC is that it **normalizes** the quartet graph to account for “artificial taxa,” which are introduced so that solutions on subproblems can be combined during the divide phase (this framework naturally accommodates multi-labeled trees as input). To normalize the graph efficiently, TQMC requires the number of leaves in a gene tree labeled by a taxon to be indicative of the number of quartets that taxon participates in. This relationship breaks down when excluding DQs. Therefore, TQMC-pro does not support graph normalization; however, it did not impact accuracy in our study.

## Empirical study

Next, we describe an empirical study evaluating the exclusion-only objective function implemented within TQMC-pro. Software versions and exact commands are available in the Supplement. Data are available on Box (https://umd.box.com/v/tqmc-pro-data) and will be moved to a repository pending acceptance.

### Simulation Study

To evaluate TQMC-pro, data were simulated under the DLCoal model with 25 taxa (and an outgroup) and 2000 gene trees using the same protocol as the A-pro study [Zhang et al., 2020]. Re-simulation was necessary to save the true duplication/loss events, which were needed to evaluate tagging accuracy. Full simulation parameters are provided in Table S1 of the Supplement and summarized below.

The first experiment varied the gene duplication rate as well as the effective population size, which in turn **varied the ILS level** (loss rate of zero). Data were simulated with four duplication levels (0, 1, 2, 5) and four ILS levels (0, 0.2, 0.5, 0.7), where duplication level was evaluated as the mean number of gene copies per species minus one and ILS level was evaluated as the mean RF distance between true locus and gene tree pairs for the dup=0 condition. In total, there were 16 model conditions, each with 25 replicates.

The second experiment varied **gene tree estimation error (GTEE)**. Molecular sequences of 500bp were simulated under the GTR model for the first 10 replicates of each condition described above. After, gene trees were estimated from the first 100bp and 500bp under the GTR model using IQ-TREE [Minh et al., 2020]. Gene tree estimation error (GTEE) was evaluated as the mean RF distance between the true and estimated gene trees. For 100bp and 500bp, GTEE was around 0.5 and 0.2, respectively.

The third experiment varied the duplication rate *λ* as well as **loss/duplication rate ratio** *µ*/*λ* (effective population size was set to the same value as the 0.5 ILS level). Data were simulated with four duplication levels (0, 1, 2, 5) and three ratios (0, 0.5, 1) (the simulator did not allow *µ* > *λ*; see the Supplement for details). In total, there were 9 model conditions, each with 25 replicates.

### Evaluation Metrics

The accuracy of gene tree tagging was evaluated for each pair of gene copies (i.e., leaves). Specifically, TPs are paralogs in both the true and A-pro taggings, FPs are orthologs in the true tagging and paralogs in the A-pro tagging, and FNs are paralogs in the true tagging but orthologs in the A-pro tagging. Precision and recall were computed for each model condition taking the first 100 gene trees of the 10 replicates. Species tree error was evaluated as the normalized RF distance between the true and estimated species trees; it corresponds to the fraction of FN/FP branches, as both trees are binary.

### Methods

TQMC-pro was evaluated against A-pro and standard quartet-based species tree methods: A-multi and TQMC, which is similar to A-multi. All methods were executed given the first 250, 500, 1000, or 2000 gene trees per replicate data set. We ran TQMC in default mode, which uses n2 graph normalization, as well as without normalization (n0) to match TQMC-pro. For each data set, the true duplication events were used to tag the true rooted gene trees based on our definition of correct tagging. Additionally, true and estimated gene trees were rooted and tagged using A-pro. These two sets of trees were given as inputs to TQMC-pro. To our knowledge, A-pro does not take tagged gene trees as input and thus performed rooting and tagging internally.

### Plant re-analysis

Lastly, we re-analyzed the 1kp plant data set [Wickett et al., 2014] with 83 taxa and 9,237 gene family trees using A-pro, TQMC-pro (with A-pro tagged gene trees), and A-multi. Trees were compared based on their quartet scores computed with both A-pro and A-multi. We also evaluated differences against the ASTRAL tree (80 taxa) estimated from single-copy genes, reporting support (local posterior probability) computed with either A-pro or A-multi on differing branches.

## Empirical Results and Discussion

We now describe the results of simulations and plant re-analysis.

### Varying Duplication and ILS Levels

We first looked at interactions between varying the duplication and ILS levels, holding the loss rate at 0. The number of duplication vertices in the gene trees increases with ILS level, jumping from ∼23 to ∼31 for the highest duplication level (5), as suggested by our example (Fig. 1). Tagging accuracy increased with duplication level (although so did the number of pairwise comparisons) but decreased with ILS level. Nevertheless, precision was greater than 0.75 and recall was greater than 0.8, across all model conditions (Fig. 2A). Perhaps as a result, there was little to no difference in species tree error between TQMC-pro given the true tagging or A-pro’s rooting/tagging (Fig. 2B). Likewise, TQMC-pro and A-pro achieved similar error rates across all model conditions and outperformed the “multi” methods, with differences most pronounced for small numbers of genes, higher duplication levels, and high ILS levels. The performance of “pro” methods tended to improve with increasing duplication level, whereas the performance of “multi” methods deteriorated. As an example, at the highest ILS level (0.7) and smallest number of genes (250), the pro method error *decreased* from ∼0.1 to <0.05 when going from duplication level 0 to 5, whereas the multi method error *increased* from ∼0.1 to >0.15 (A-multi and TQMC-n0) or stayed the same (TQMC-n2). One possible explanation is that duplications produce more conflicting quartets for the multi methods but not the pro methods (additionally, quartets impacted by deep coalescence are more likely to be duplication quartets). It is also worth noting that the ILS level decreased from 0.71 for dup=0 to 0.62 for dup=5 in this example, perhaps due to the added constraints of the bounded coalescent, so the model condition became less challenging (Table S2).

### Varying GTEE

Next, we evaluated the impact of gene tree estimation error. Tagging precision decreased as GTEE increased, whereas recall stayed largely the same. This suggests an increase in FPs (Fig. 2A,C,D), and relatedly, the number of vertices tagged as duplications by A-pro increased with GTEE (Table S2). Likewise, species tree error typically increased as GTEE increased, when holding model conditions constant, especially for A-multi and TQMC-n0 (Figs. 2E and S1), although there were exceptions at the highest ILS levels (Figs. S2–S3). In the context of GTEE, increasing the duplication level sometimes decreased species tree error, even for the multi-methods (especially for duplication levels 1-2 and small numbers of gene trees). One possible explanation is that the greater number of quartets available from duplications has an advantage when there is noise.

### Varying Loss/Duplication Rate Ratio

Next, we evaluated the impact of varying the loss/duplication rate ratio, holding the effective population size constant (ILS level of 0.5). Losses had little impact on either tagging accuracy (Fig. S4) or species tree error (Fig. S5). However, increasing the loss rate also lowered the amount of ILS, especially as the duplication rate increased (Table S3), likely related to the bounded coalescent.

### Plant Re-analysis

Lastly, we re-analyzed the 1kp plant data set. Both ASTRAL-pro and TQMC-pro produced similar trees to the published ASTRAL tree on single-copy genes, differing by just 5 and 4 branches out of 77, respectively, and recovering all major clades (Fig. 3 and Fig. S6). They differed from each other by just 4 branches, often with lower branch support. In contrast, A-multi differed from the ASTRAL tree by 58 branches and failed to recover the major clades. The inter-mixing of these clades often led to branch support values of 0. To rule out the possibility that A-multi had just failed to find a good solution to its optimization problem, we computed the A-multi score for both the A-pro and TQMC-pro trees. We found that the A-multi tree was indeed the highest scoring, suggesting that the problem is due to its objective function not excluding duplication quartets.

**Fig. 3.**
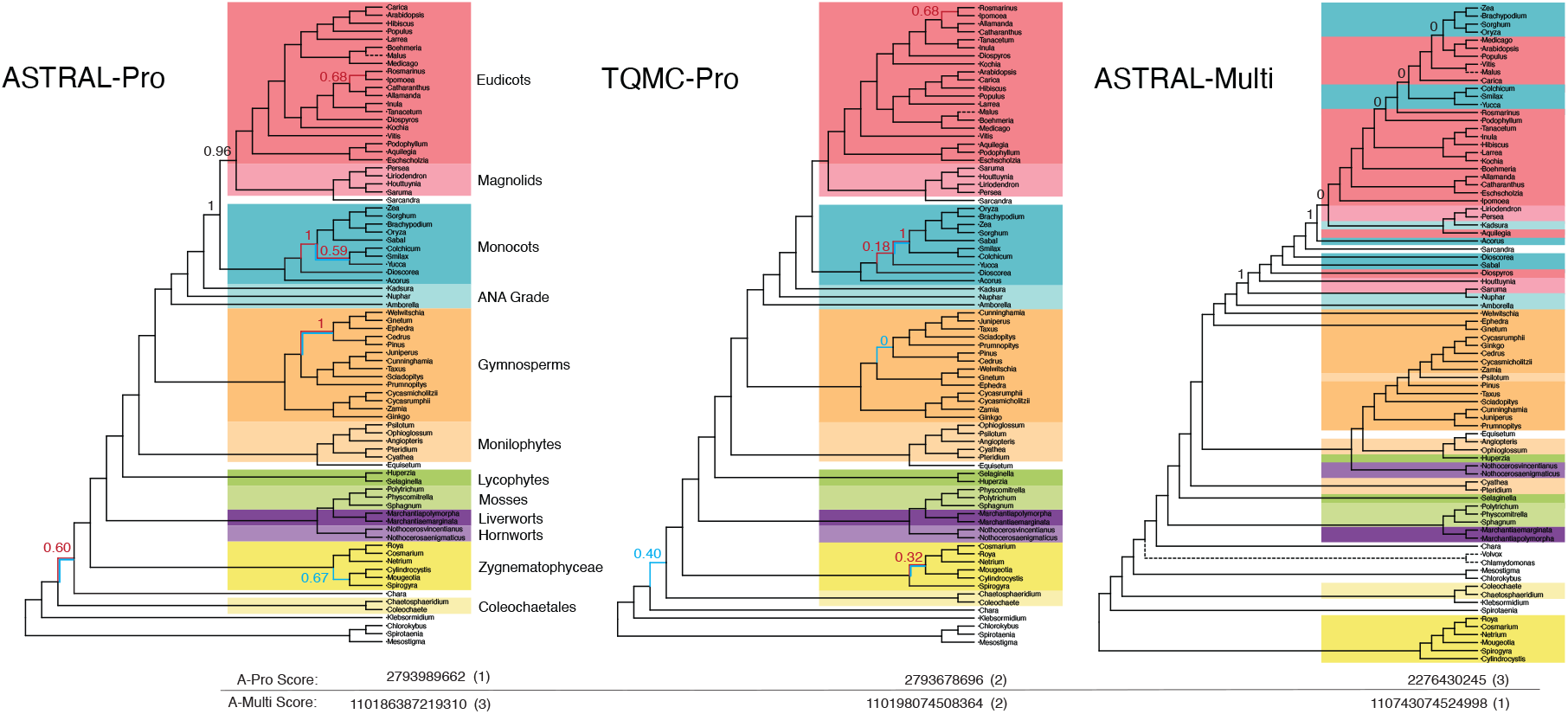
Species trees estimated by A-Pro, TQMC-pro, and A-Multi on the multi-copy 1kp plant data set. All trees were rooted at (*Uronema*, (*V olvox, Chlamydomonas*)), not shown. Dashed lines indicate the species was missing from the single-copy ASTRAL tree. The A-pro tree and TQMC-pro tree differed from the single-copy ASTRAL tree by 5 and 4 branches, respectively, shown in red; they differed from each other by four branches; shown in blue. Branch support (range 0 to 1) computed with A-pro is indicated on these branches as well as on the clades for Monocots and Eudicots+Magnolids, which are split in the A-multi tree. Branch support computed by A-multi is shown for select branches where these clades intermix. Quartet scores are shown at the bottom.

## Conclusions

In this work, we presented a definition of correct gene tree tagging and studied it from a theoretical and empirical perspective. For the former, we found that under our definition of correct tagging only gene copies that descend from the same locus lineage will be SQs (simplifying the number of locus (also called root) lineage scenarios that need to be considered, compared to Markin and Eulenstein [2021], Hill et al. [2022]). Unfortunately, our definition still results in a breakdown of a straightforward exchangeability argument in the case of deep coalescence; thus, more delicate argumentation is necessary, and whether or not (an exclusion only version of) A-pro is consistent is an open question. However, we showed empirically that DQ exclusion within TQMC (called TQMC-pro) achieves similar performance to A-pro on real and simulated data sets. Additionally, we observed that although tagging accuracy decreased with the amount of ILS and estimation error, species tree accuracy stayed high. Errors in tagging (evaluated pairwise across leaves) does not necessarily translate to errors in DQ/SQ classification. In particular, false positives (classifying a speciation vertex as a duplication vertex) would flip SQs to DQs, which may not impact species tree accuracy, especially if those SQs were impacted by ILS, provided there was sufficient phylogenetic signal.

## Acknowledgements

The authors thank Yunheng Han for helpful discussions about the TREE-QMC code base. This material is based upon work supported by the U. S. National Science Foundation (NSF) under Grant Number 2441458 (to EKM) as well as the State of Maryland. AM was supported in part by the USDA-ARS (ARS project number 5030-32000-231-000D) and USDA-APHIS (ARS project number 5030-32000-231-111-I). This work was initiated at the NSF-funded Institute for Computational and Experimental Research in Mathematics. The funders had no role in study design, data collection, and interpretation, or the decision to submit the work for publication. Mention of trade names or commercial products in this article is solely for the purpose of providing specific information and does not imply recommendation or endorsement by the USDA. USDA is an equal opportunity provider and employer.

