## supplement for "On correctness of gene tree tagging under a unified model of gene duplication, loss, and coalescence"

April 10, 2026

**4 Supplemental Results** 12

**References** 18

### 1 Supplemental Methods

#### 1.1 TREE-QMC-Pro

The main contribution of TREE-QMC is it constructs the quartet graph for each subproblem directly from the input gene trees, rather than enumerating all quartets. Consider the following. In a traversal of an arbitrarily rooted gene tree, each internal node contributes good and bad edges to the graph based on the quartets that can be formed around the node. Suppose we wish to compute the number of bad edges between species $X$ and $Y$ *around* node $t$, where $X$ is below $t.l$ and $Y$ is below $t.r$; this quantity is denoted $\mathbb{B}_t(X, Y)$ (note that swapping $X, Y$ reverses the relationship). Let $t.l$ and $t.r$ denote the left and right child of $t$, respectively. Then, we must count all ways of selecting two leaves $x, y$ labeled $X, Y$ plus two leaves $z, w$ with other labels $Z, W$ ($X \neq Y \neq Z \neq W$) such that the resulting quartet has $X$ and $Y$ as siblings. There are three ways this can occur:

1. $x$ forms the $(x,(z,w))$ triplet with $z$ and $w$ below $t.l$ and $y$ is below $t.r$
2. $x$ is below $t.l$ and $y$ forms the $(y,(z,w))$ triplet below $t.r$ (symmetric to case 1)
3. $x$ is below $t.l$, $y$ is below $t.r$, and $z, w$ are selected from above $t$

Now suppose we wish to compute the number of good edges between species $X$ and $Y$ around node $t$, denoted $\mathbb{G}_t(X, Y)$. Now, we count all the ways of selecting two leaves $x, y$ labeled $X, Y$ and two other leaves $z, w$ labeled $Z, W$ ($X \neq Y \neq Z \neq W$) such that the resulting quartet has $X$ and $Y$ on opposite sides of the internal branch. We have the following cases (note that selection of $Z$ and $W$ is exchangeable):

1. $x$ forms the triplet $(w,(z,x))$ below $t.l$ and $y$ is below $t.r$
2. $x$ is below $t.l$ and $y$ forms the triplet $(w,(z,y))$ below $t.r$ of $t$ (symmetric to case 1)
3. $x$ forms a doublet with $z$ below $t.l$, $y$ is below $t.r$, and $w$ is above $t$
4. $x$ is below $t.l$, $y$ forms a doublet with $z$ below $t.r$, and $w$ is above $t$ (symmetric to case 3)
5. $x$ form a doublet with $z$ below $t.l$ and $y$ forms a doublet with $w$ below $t.r$

where "form a doublet below $t$" means their MRCA is a descendant of $t$.

Several quantities above are used across multiple cases. Therefore, we can precompute these values, which we called "auxillary values," to improve the efficiency of computing bad and good edges. The corresponding equations for the auxillary values computed by TREE-QMC are:

1. **Singlet below:** number of times species $X$ appears below node $t$ (computed in postorder traversal)

$$\mathrm{w}^1_X(t) = \begin{cases} 1 & \text{if } t \text{ is a leaf} \\ \mathrm{w}^1_X(t.l) + \mathrm{w}^1_X(t.r) & \text{otherwise} \end{cases} \tag{1}$$

2. **Singlet above:** number of times species $X$ appears above node $t$ (computed in preorder traversal)

$$\overline{\mathrm{w}}^1_X(t) = \begin{cases} 0 & \text{if } t \text{ is the root} \\ \overline{\mathrm{w}}^1_X(t.p) + \mathrm{w}^1_X(t.s) & \text{otherwise} \end{cases} \tag{2}$$

3. **Doublet below:** number of times species $X$ and $Y$ can form a doublet below node $t$ (postorder)

$$\mathrm{w}^2_{X,Y}(t) = \mathrm{w}^2_{Y,X}(t) = \begin{cases} 0 & \text{if } t \text{ is a leaf} \\ \mathrm{w}^2_{X,Y}(t.l) + \mathrm{w}^2_{X,Y}(t.r) + \mathrm{w}^1_X(t.l) + \mathrm{w}^1_Y(t.r) & \text{otherwise} \\ +\mathrm{w}^1_Y(t.l) + \mathrm{w}^1_X(t.r) \end{cases} \tag{3}$$

4. **Doublet above:** number of times species $X$ and $Y$ can form a doublet above node $t$ (preorder)

$$\overline{\mathrm{w}}^2_{X,Y}(t) = \overline{\mathrm{w}}^2_{Y,X}(t) = \begin{cases} 0 & \text{if } t \text{ is the root} \\ \overline{\mathrm{w}}^2_{X,Y}(t.p) + \mathrm{w}^2_{X,Y}(t.s) + \overline{\mathrm{w}}^1_X(t.p) \cdot \mathrm{w}^1_Y(t.s) & \text{otherwise} \\ +\mathrm{w}^1_X(t.s) \cdot \overline{\mathrm{w}}^1_Y(t.p) \end{cases} \tag{4}$$

5. **Bad triplet** number of times triplet $(X,(Z,W))$ occurs below node $t$ for all possible $Z, W$ (postorder)

$$\mathrm{w}^{x|z,w}_{X,Y}(t) = \begin{cases} 0 & \text{if } t \text{ is a leaf} \\ \mathrm{w}^{x|z,w}_{X,Y}(t.l) + \mathrm{w}^{x|z,w}_{X,Y}(t.r) & \text{otherwise} \\ +\mathrm{w}^1_X(t.l) \cdot \sum_{\substack{Z,W \in \mathcal{S}\cup\mathcal{A} \\ Z\neq W\neq X\neq Y}} \mathrm{w}^2_{Z,W}(t.r) \\ +\mathrm{w}^1_X(t.r) \cdot \sum_{\substack{Z,W \in \mathcal{S}\cup\mathcal{A} \\ Z\neq W\neq X\neq Y}} \mathrm{w}^2_{Z,W}(t.l) \end{cases} \tag{5}$$

6. **Good triplet:** number of times triplet $(W,(Z,X))$ occurs below node $t$ for all possible $Z, W$ (postorder)

$$\mathrm{w}^{x,z|w}_{X,Y}(t) = \begin{cases} 0 & \text{if } t \text{ is a leaf} \\ \mathrm{w}^{x,z|w}_{X,Y}(t.l) + \mathrm{w}^{x,z|w}_{X,Y}(t.r) & \text{otherwise} \\ + \sum_{\substack{Z\in\mathcal{S}\cup\mathcal{A} \\ Z\neq X,Y}} \sum_{\substack{W\in\mathcal{S}\cup\mathcal{A} \\ W\neq X,Y,Z}} \mathrm{w}^2_{X,Z}(t.l)\mathrm{w}^1_W(t.r) \\ + \sum_{\substack{Z\in\mathcal{S}\cup\mathcal{A} \\ Z\neq X,Y}} \sum_{\substack{W\in\mathcal{S}\cup\mathcal{A} \\ W\neq X,Y,Z}} \mathrm{w}^2_{X,Z}(t.r)\mathrm{w}^1_W(t.l) \end{cases} \tag{6}$$

After auxiliary values, the bad and good edges at node $t$ can be computed via postorder traversals as:

$$\Delta\mathbb{B}_t(X,Y) = \begin{cases} \mathrm{w}^1_Y(t.r) \cdot \mathrm{w}^{x|z,w}_{X,Y}(t.l) \\ +\mathrm{w}^1_X(t.l) \cdot \mathrm{w}^1_Y(t.r) \cdot \sum_{\substack{Z,W\in\mathcal{S}\cup\mathcal{A} \\ Z\neq W\neq X\neq Y}} \overline{\mathrm{w}}^2_{Z,W}(t) \\ +\mathrm{w}^1_X(t.l) \cdot \mathrm{w}^{x|z,w}_{Y,X}(t.r) \end{cases} \tag{7}$$

$$\Delta\mathbb{G}_t(X,Y) = \begin{cases} \mathrm{w}^1_Y(t.r) \cdot \mathrm{w}^{x,z|w}_{X,Y}(t.l) \\ +\mathrm{w}^1_Y(t.r) \cdot \sum_{\substack{Z\in\mathcal{S}\cup\mathcal{A} \\ Z\neq X,Y}} \sum_{\substack{W\in\mathcal{S}\cup\mathcal{A} \\ W\neq X,Y,Z}} \mathrm{w}^2_{X,Z}(t.l) \cdot \overline{\mathrm{w}}^1_W(t) \\ + \sum_{\substack{Z\in\mathcal{S}\cup\mathcal{A} \\ Z\neq X,Y}} \sum_{\substack{W\in\mathcal{S}\cup\mathcal{A} \\ W\neq X,Y,Z}} \mathrm{w}^2_{X,Z}(t.l) \cdot \mathrm{w}^2_{Y,W}(t.r) \\ +\mathrm{w}^1_X(t.l) \cdot \sum_{\substack{Z\in\mathcal{S}\cup\mathcal{A} \\ Z\neq X,Y}} \sum_{\substack{W\in\mathcal{S}\cup\mathcal{A} \\ W\neq X,Y,Z}} \mathrm{w}^2_{Y,Z}(t.r) \cdot \overline{\mathrm{w}}^1_W(t) \\ +\mathrm{w}^1_X(t.l) \cdot \mathrm{w}^{x,z|w}_{Y,X}(t.r) \end{cases} \tag{8}$$

Weighted TREE-QMC uses the quartet weighting introduced by [9]. This scheme weights quartets based on the terminal branch lengths and internal node support. We let $W_l(q)$ be the weight contribution of the terminal branch lengths of the quartet $q$ and $W_s(q)$ be the weight contribution of the internal branch support. We define $W_l(q)$ and $W_s(q)$ as follows:

$$W_l(q) = \exp\left( \sum_{\substack{e\in x,y\to u, \\ z,w\to v}} -l(e) \right) \tag{9}$$

$$W_s(q) = 1 - \prod_{e \in u \to v} (1 - s(e)) \tag{10}$$

where $l(e)$ denotes the length of branch $e$ in the gene tree, $s(e)$ denotes the support value of branch $e$, $u$ and $v$ denote the anchor vertices of quartet $q = xy|zw$ where $u$ is closer to siblings $x, y$ and $v$ is closer to $z, w$. We let $u \to v$ be the set of edges on the path from $u$ to $v$ (similarly for $x, y \to u$ and $z, w \to v$). The total weight of the quartet is then $W_h(q) = W_l(q) \cdot W_s(q)$. Introducing this weighting scheme gives us the following updated auxiliary value equations:

1. **Singlet below:**

$$\mathrm{w}^1_X(t) = \begin{cases} I(t) & \text{if } t \text{ is a leaf} \\ \exp(-\mathrm{l}(t.l)) \cdot \mathrm{w}^1_X(t.l) + \exp(-\mathrm{l}(t.r)) \cdot \mathrm{w}^1_X(t.r) & \text{otherwise} \end{cases} \tag{11}$$

where $I(t)$ is the importance value of the taxon labeling $t$. This importance value is set to normalize the graph; see [2, 3] for details. It is always set to one when normalization is not used, as is the case for TREE-QMC-pro.

2. **Singlet above:**

$$\overline{\mathrm{w}}^1_X(t) = \begin{cases} 0 & \text{if } t \text{ is the root} \\ \exp(-\mathrm{l}(t)) \cdot (\overline{\mathrm{w}}^1_X(t.p) + \exp(-\mathrm{l}(t.s)) \cdot \mathrm{w}^1_X(t.s)) & \text{otherwise} \end{cases} \tag{12}$$

3. **Doublet below:**

$$\mathrm{w}^2_{X,Y}(t) = \mathrm{w}^2_{Y,X}(t) = \begin{cases} 0 & \text{if } t \text{ is a leaf} \\ (1 - \mathrm{s}(t.l)) \cdot \mathrm{w}^2_{X,Y}(t.l) + (1 - \mathrm{s}(t.r)) \cdot \mathrm{w}^2_{X,Y}(t.r) & \text{otherwise} \\ +\exp(-\mathrm{l}(t.l)) \cdot \mathrm{w}^1_X(t.l) + \exp(-\mathrm{l}(t.r)) \cdot \mathrm{w}^1_Y(t.r) & \\ +\exp(-\mathrm{l}(t.l)) \cdot \mathrm{w}^1_Y(t.l) + \exp(-\mathrm{l}(t.r)) \cdot \mathrm{w}^1_X(t.r) & \end{cases} \tag{13}$$

4. **Doublet above:**

$$\overline{\mathrm{w}}^2_{X,Y}(t) = \overline{\mathrm{w}}^2_{Y,X}(t) = \begin{cases} 0 & \text{if } t \text{ is the root} \\ (1 - \mathrm{s}(t)) \cdot \overline{\mathrm{w}}^2_{X,Y}(t.p) & \text{otherwise} \\ +(1 - \mathrm{s}(t)) \cdot (1 - \mathrm{s}(t.s)) \cdot \mathrm{w}^2_{X,Y}(t.s) & \\ +(1 - \mathrm{s}(t)) \cdot \exp(-\mathrm{l}(t.s)) \cdot \overline{\mathrm{w}}^1_X(t.p) \cdot \mathrm{w}^1_Y(t.s) & \\ +(1 - \mathrm{s}(t)) \cdot \exp(-\mathrm{l}(t.s)) \cdot \mathrm{w}^1_X(t.s) \cdot \overline{\mathrm{w}}^1_Y(t.p) & \end{cases} \tag{14}$$

5. **Bad triplet:**

$$\mathrm{w}^{x|z,w}_{X,Y}(t) = \begin{cases} 0 & \text{if } t \text{ is a leaf} \\ \exp(-\mathrm{l}(t.l)) \cdot \mathrm{w}^{x|z,w}_{X,Y}(t.l) + \exp(-\mathrm{l}(t.r)) \cdot \mathrm{w}^{x|z,w}_{X,Y}(t.r) & \text{otherwise} \\ +\exp(-\mathrm{l}(t.l)) \cdot \mathrm{w}^1_X(t.l) \cdot (1 - \mathrm{s}(t.r)) \cdot \sum_{\substack{Z,W \in \mathcal{S} \cup \mathcal{A} \\ Z \neq W \neq X \neq Y}} \mathrm{w}^2_{Z,W}(t.r) & \\ +\exp(-\mathrm{l}(t.r)) \cdot \mathrm{w}^1_X(t.r) \cdot (1 - \mathrm{s}(t.l)) \cdot \sum_{\substack{Z,W \in \mathcal{S} \cup \mathcal{A} \\ Z \neq W \neq X \neq Y}} \mathrm{w}^2_{Z,W}(t.l) & \end{cases} \tag{15}$$

6. **Good triplet:**

$$\mathrm{w}^{x,z|w}_{X,Y}(t) = \begin{cases} 0 & \text{if } t \text{ is a leaf} \\ \exp(-\mathrm{l}(t.l)) \cdot \mathrm{w}^{x,z|w}_{X,Y}(t.l) & \text{otherwise} \\ +\exp(-\mathrm{l}(t.r)) \cdot \mathrm{w}^{x,z|w}_{X,Y}(t.r) & \\ +(1 - \mathrm{s}(t.l)) \cdot \exp(-\mathrm{l}(t.r)) \cdot \sum_{\substack{Z \in \mathcal{S} \cup \mathcal{A} \\ Z \neq X,Y}} \sum_{\substack{W \in \mathcal{S} \cup \mathcal{A} \\ W \neq X,Y,Z}} \mathrm{w}^2_{X,Z}(t.l) \mathrm{w}^1_W(t.r) & \\ +(1 - \mathrm{s}(t.r)) \cdot \exp(-\mathrm{l}(t.l)) \cdot \sum_{\substack{Z \in \mathcal{S} \cup \mathcal{A} \\ Z \neq X,Y}} \sum_{\substack{W \in \mathcal{S} \cup \mathcal{A} \\ W \neq X,Y,Z}} \mathrm{w}^2_{X,Z}(t.r) \mathrm{w}^1_W(t.l) & \end{cases} \tag{16}$$

The intuition is that for whenever the value being computed is on an internal edge, the support value must contribute to the score and whenever the value is contributing to a terminal branch in the quartet, the branch length needs to be included. Note that these equations can also be used in the unweighted case, where all quarets have weight 1, by setting $l(e) = 0$ for all edges and $s(e) = 1$ for all edges. We only explore the unweighted case in our study but because we update these recurrences, we get quartet weighting for free.

Lastly, we need to update these equations to only count speciation quartets (SQs). We can assume that the tree has been tagged with duplications following the algorithm in ASTRAL-Pro. To exclude the contribution of duplication quartets, we need to consider all ways that a duplication quartet can be formed in the input tree. We can consider again the ways taxa $X$ and $Y$ can contribute a bad edge to the quartet graph. In case where $X$ forms the $(X,(Z,W))$ triplet with $Z$ and $W$ below $t.l$ and $Y$ is below $t.r$, there are two nodes that must both be tagged as speciation for the quartet to be an SQ and thus counted: (1) the MRCA of $X, W, Z$ and (2) the MRCA of $X, Y, W, Z$. If either of these nodes are tagged as duplication (dup), the quartet is a DQ and should be excluded. We access the node in (1) when computing the triplet auxiliary value and node (2) in the bad edge computation. Using this approach for the other cases for both good and bad edges, we find that checking for duplication tagged nodes must be done in the singlet above, doublet above, good triplet, and bad triplet auxiliary value equations as well as in the equations for calculating the good and bad edges. The updated equations are:

1. Singlet above

$$\overline{\mathrm{w}}^1_X(t) = \begin{cases} 0 & \text{if } t \text{ is the root} \\ \exp(-\mathrm{l}(t)) \cdot \overline{\mathrm{w}}^1_X(t.p) & \text{if } t.p \text{ is tagged dup} \\ \exp(-\mathrm{l}(t)) \cdot (\overline{\mathrm{w}}^1_X(t.p) + \exp(-\mathrm{l}(t.s)) \cdot \mathrm{w}^1_X(t.s)) & \text{otherwise} \end{cases} \tag{17}$$

2. Doublet above

$$\overline{\mathrm{w}}^2_{X,Y}(t) = \overline{\mathrm{w}}^2_{Y,X}(t) = \begin{cases} 0 & \text{if } t \text{ is the root} \\ (1-\mathrm{s}(t)) \cdot \overline{\mathrm{w}}^2_{X,Y}(t.p) & \text{if } t.p \text{ is tagged dup} \\ (1-\mathrm{s}(t)) \cdot \overline{\mathrm{w}}^2_{X,Y}(t.p) + (1-\mathrm{s}(t)) \cdot (1-\mathrm{s}(t.s)) \cdot \mathrm{w}^2_{X,Y}(t.s) & \text{otherwise} \\ +(1-\mathrm{s}(t)) \cdot \exp(-\mathrm{l}(t.s)) \cdot \overline{\mathrm{w}}^1_X(t.p) \cdot \mathrm{w}^1_Y(t.s) & \\ +(1-\mathrm{s}(t)) \cdot \exp(-\mathrm{l}(t.s)) \cdot \mathrm{w}^1_X(t.s) \cdot \overline{\mathrm{w}}^1_Y(t.p) & \end{cases} \tag{18}$$

3. Bad triplet

$$\mathrm{w}^{x|z,w}_{X,Y}(t) = \begin{cases} 0 & \text{if } t \text{ is a leaf} \\ \exp(-\mathrm{l}(t.l)) \cdot \mathrm{w}^{x|z,w}_{X,Y}(t.l) + \exp(-\mathrm{l}(t.r)) \cdot \mathrm{w}^{x|z,w}_{X,Y}(t.r) & \text{if } t \text{ is tagged dup} \\ \exp(-\mathrm{l}(t.l)) \cdot \mathrm{w}^{x|z,w}_{X,Y}(t.l) + \exp(-\mathrm{l}(t.r)) \cdot \mathrm{w}^{x|z,w}_{X,Y}(t.r) & \text{otherwise} \\ +\exp(-\mathrm{l}(t.l)) \cdot \mathrm{w}^1_X(t.l) \cdot (1-\mathrm{s}(t.r)) \cdot \sum_{\substack{Z,W \in \mathcal{S} \cup \mathcal{A} \\ Z \neq W \neq X \neq Y}} \mathrm{w}^2_{Z,W}(t.r) & \\ +\exp(-\mathrm{l}(t.r)) \cdot \mathrm{w}^1_X(t.r) \cdot (1-\mathrm{s}(t.l)) \cdot \sum_{\substack{Z,W \in \mathcal{S} \cup \mathcal{A} \\ Z \neq W \neq X \neq Y}} \mathrm{w}^2_{Z,W}(t.l) & \end{cases} \tag{19}$$

4. Good triplet

$$\mathrm{w}^{x,z|w}_{X,Y}(t) = \begin{cases} 0 & \text{if } t \text{ is a leaf in } T \\ \exp(-\mathrm{l}(t.l)) \cdot \mathrm{w}^{x,z|w}_{X,Y}(t.l) + \exp(-\mathrm{l}(t.r)) \cdot \mathrm{w}^{x,z|w}_{X,Y}(t.r) & \text{if } t \text{ is a duplication node} \\ \exp(-\mathrm{l}(t.l)) \cdot \mathrm{w}^{x,z|w}_{X,Y}(t.l) + \exp(-\mathrm{l}(t.r)) \cdot \mathrm{w}^{x,z|w}_{X,Y}(t.r) & \text{otherwise} \\ +(1-\mathrm{s}(t.l)) \cdot \exp(-\mathrm{l}(t.r)) \cdot \sum_{\substack{Z \in \mathcal{S} \cup \mathcal{A} \\ Z \neq X,Y}} \sum_{\substack{W \in \mathcal{S} \cup \mathcal{A} \\ W \neq X,Y,Z}} \mathrm{w}^2_{X,Z}(t.l) \mathrm{w}^1_W(t.r) & \\ +(1-\mathrm{s}(t.r)) \cdot \exp(-\mathrm{l}(t.l)) \cdot \sum_{\substack{Z \in \mathcal{S} \cup \mathcal{A} \\ Z \neq X,Y}} \sum_{\substack{W \in \mathcal{S} \cup \mathcal{A} \\ W \neq X,Y,Z}} \mathrm{w}^2_{X,Z}(t.r) \mathrm{w}^1_W(t.l) & \end{cases} \tag{20}$$

5. Good edges

$$
\Delta\mathbb{G}_t(X,Y) = \begin{cases} 0 & \text{if } t \text{ is tagged dup} \\ \mathrm{w}^1_Y(t.r)\cdot\exp(-\mathrm{l}(t.r))\cdot \mathrm{w}^{x,z|w}_{X,Y}(t.l)\cdot\exp(-\mathrm{l}(t.l)) & \text{otherwise} \\ +\exp(-\mathrm{l}(t.r))\cdot \mathrm{w}^1_Y(t.r)\cdot(1-\mathrm{s}(t.l))\cdot \sum_{\substack{Z\in\mathcal{S}\cup\mathcal{A}\\ Z\neq X,Y}}\sum_{\substack{W\in\mathcal{S}\cup\mathcal{A}\\ W\neq X,Y,Z}} \mathrm{w}^2_{X,Z}(t.l)\cdot\overline{\mathrm{w}}^1_W(t) & \\ +(1-\mathrm{s}(t.l))\cdot(1-\mathrm{s}(t.r))\cdot \sum_{\substack{Z\in\mathcal{S}\cup\mathcal{A}\\ Z\neq X,Y}}\sum_{\substack{W\in\mathcal{S}\cup\mathcal{A}\\ W\neq X,Y,Z}} \mathrm{w}^2_{X,Z}(t.l)\cdot \mathrm{w}^2_{Y,W}(t.r) & \\ +\exp(-\mathrm{l}(t.l))\cdot \mathrm{w}^1_X(t.l)\cdot(1-\mathrm{s}(t.r))\cdot \sum_{\substack{Z\in\mathcal{S}\cup\mathcal{A}\\ Z\neq X,Y}}\sum_{\substack{W\in\mathcal{S}\cup\mathcal{A}\\ W\neq X,Y,Z}} \mathrm{w}^2_{Y,Z}(t.r)\cdot\overline{\mathrm{w}}^1_W(t) & \\ +\mathrm{w}^1_X(t.l)\cdot\exp(-\mathrm{l}(t.l))\cdot \mathrm{w}^{x,z|w}_{Y,X}(t.r)\cdot\exp(-\mathrm{l}(t.r)) & \end{cases} \tag{21}
$$

6. Bad edges

$$
\Delta\mathbb{B}_t(X,Y) = \begin{cases} \exp(-\mathrm{l}(t.l))\cdot \mathrm{w}^1_X(t.l)\cdot\exp(-\mathrm{l}(t.r))\cdot \mathrm{w}^1_Y(t.r)\cdot \sum_{\substack{Z,W\in\mathcal{S}\cup\mathcal{A}\\ Z\neq W\neq X\neq Y}} \overline{\mathrm{w}}^2_{Z,W}(t) & \text{if } t \text{ is tagged dup} \\ \mathrm{w}^1_Y(t.r)\cdot\exp(-\mathrm{l}(t.r))\cdot \mathrm{w}^{x|z,w}_{X,Y}(t.l)\cdot\exp(-\mathrm{l}(t.l)) & \text{otherwise} \\ +\exp(-\mathrm{l}(t.l))\cdot \mathrm{w}^1_X(t.l)\cdot\exp(-\mathrm{l}(t.r))\cdot \mathrm{w}^1_Y(t.r)\cdot \sum_{\substack{Z,W\in\mathcal{S}\cup\mathcal{A}\\ Z\neq W\neq X\neq Y}} \overline{\mathrm{w}}^2_{Z,W}(t) & \\ +\mathrm{w}^1_X(t.l)\cdot\exp(-\mathrm{l}(t.l))\cdot \mathrm{w}^{x|z,w}_{Y,X}(t.r)\cdot\exp(-\mathrm{l}(t.r)) & \end{cases} \tag{22}
$$

#### 1.2 Simulated Data

**Species trees and gene trees.** In our study, we simulate datasets based on the protocol from the ASTRAL-Pro study [10] with varying the effective population size as well as the duplication and loss rates. Species trees and gene trees were simulated using Simphy. To obtain correct node-mapping information between species tree and gene tree nodes, we compiled SimPhy from the postorderSptree branch, available at https://github.com/adamallo/SimPhy/tree/postorderSptree (commit 4d1063e). SimPhy v1.0 was then run using the following example command:

```
simphy \
    -sl f:25 \
    -rs 25 \
    -rl f:2000 \
    -rg 1 \
    -sb f:0.000000005 \
    -sd f:0 \
    -st ln:21.25,0.2 \
    -so f:1 \
    -si f:1 \
    -sp f:<global population sizes> \
    -su ln:-21.9,0.1 \
    -hh f:1 \
    -hs ln:1.5,1 \
    -hl ln:1.551533,0.6931472 \
    -hg ln:1.5,1 \
    -cs 9644 \
    -v 1 \
    -ot 0 \
    -op 1 \
    -om 1 \
    -ol 1 \
    -lb f:<duplication rate> \
    -ld f:<loss rate> \
    -lt f:0
```

The full simulation parameters set is given in Table S1. Note that SimPhy enforces a constraint requiring loss rate to less than or equal to the duplication rate. When this is violated, SimPhy terminates with the following message: `Settings error: Improper value sampling the parameter -Ld, Locus tree loss rate. Please, check your sampling settings and try again.`

**DNA sequences and estimated gene trees.** For the first 10 replicates of the D1 data sets, DNA sequences of length 500 nucleotides were then simulated under the GTR+GAMMA model for each of the 2000 gene trees using INDELible v1.03 [1] with the following command:

```
indelible control.txt
```

where a new `control.txt` is generated for each species tree replicate. A representative `control.txt` file is shown below.

```
[TYPE] NUCLEOTIDE 1
[MODEL] GTR0001
 [submodel]  GTR 0.348457 0.083650 0.106221 0.047057 0.141989
 [statefreq] 0.225764 0.252186 0.334970 0.187080
 [rates] 0 1.929409 0
...
[MODEL] GTR2000
 [submodel]  GTR 0.332254 0.075895 0.076666 0.094838 0.090637
 [statefreq] 0.279294 0.175915 0.313061 0.231730
 [rates] 0 2.281270 0
[SETTINGS]
[randomseed] 2478
[fileperrep] FALSE
[TREE] T0001 <gene tree 0001 in newick string format>
...
[TREE] T2000 <gene tree 2000 in newick string format>
[PARTITIONS] T0001 [T0001 GTR0001 500]
...
[PARTITIONS] T2000 [T2000 GTR2000 500]
[EVOLVE] T0001 1 0001
...
T2000 1 2000
```

The model parameters in each control file were randomly drawn from a distribution, following the protocol used in the ASTRAL-II study [6]. Specifically, base frequencies for A,C,G,T were sampled from a Dirchlet(36,26,28,32) distribution, estimated via maximum likelihood from three large-scale empirical datasets (1KP dataset [8], Song et al Mammalian dataset [7], and Avian phylogenomics dataset [4]). GTR substitution matrices were similarly drawn from a Dirchlet(16,3,5,5,6,15) distribution. The shape parameter $\alpha$ for the gamma-distributed rates across sites was drawn from an exponential distribution with rate 1.2, discarding values below 0.1. The same seed (2478) was used in all INDELible simulations, same as in the ASTRAL-II study. To simulate different levels of gene tree estimation error (GTEE), DNA sequences were also truncated to length 100bp.

Gene trees were estimated under the GTR+GAMMA model using IQ-TREE-2 v2.4.0 [5] with the following command:

```
iqtree2 \
    -s [input alignment] \
    -m GTR+F+G4 \
    -abayes \
    -T 1
```

on sequences of length 500bp and 100bp.

Table S1: **Simulation Parameters.** Model conditions are described by (1) Dup level, which is the mean number of copies per species, (2) loss/dup rate ratio, and (3) the ILS level, which is the mean RF distance between the true locus tree and true gene tree, computed for the Dup level 0 and loss/dup ratio 0 condition.

| Argument | Description | Value |
|---|---|---|
| SL | Number of leaves | 25 |
| RS | Number of replicates | 25 |
| RL | Number of loci | 2000 |
| RG | Number of genes | 1 |
| SB | Speciation rates | 5e-9 |
| SD | Extinction rates | 0 |
| ST | Maximum tree length (time units) | LogN(21.25,0.2) |
| SO | Outgroup branch length relative to half the tree length | 1 |
| SI | Number of individuals per species | 1 |
| SU | Global substitution rate | LogN(-21.9,0.1) |
| HH | Gene-by-lineage-specific locus tree parameter | 1 |
| HS | Species-specific branch rate heterogeneity | LogN (1.5, 1) |
| HL | Locus-specific rate heterogeneity | LogN (1.551533,0.6931472) |
| HG | Gene-tree branch rate heterogeneity | LogN (1.5, 1) |
| CS | Random number generator seed | 9644 |
| OM | Tree mapping outputs | 1 |
| OL | Post-order of internal node labeling | 1 |
| LT | Horizontal gene transfer rate | 0 |
| | **D1 - Controlling duplication and ILS level (4 × 4 conditions)** | |
| SP | Global population sizes | 1e+4 (ILS=0), 4.8e+7 (ILS=0.2), 1.9e+8 (ILS=0.5), 4.7e+8 (ILS=0.7) |
| LB | Duplication rate | 0 (Dup=0), 1.9e-10 (Dup=1), 2.7e-10 (Dup=2), 4.9e-10 (Dup=5) |
| LD | Loss rate | 0 |
| | **D2 - Controlling duplication and loss rate (4 × 3 conditions)** | |
| SP | Global population sizes | 1.9e+8 |
| LB | Duplication rate | 0 (Dup=0), 1.9e-10 (Dup=1), 2.7e-10 (Dup=2), 4.9e-10 (Dup=5) |
| LD | Loss rate for ratio 0 | 0 |
| | Loss rate for ratio 0.5 | 9.5e-11 (Dup=1), 1.35e-10 (Dup=2), 2.45e-10 (Dup=5) |
| | Loss rate for ratio 1 | 1.9e-10 (Dup=1), 2.7e-10 (Dup=2), 4.9e-10 (Dup=5) |

Table S2: **D1 Summary Statistics.** All summary statistics are averaged across all 2000 gene trees and/or locus trees and then averaged across all 25 replicates for true trees and 10 replicates for estimated trees. **Dup level** is controlled by varying the duplication rate; it is then empirically evaluated as the mean number of copies per species minus one (column 9). *ILS level* is controlled by varying the effective population size; it is then empirically evaluated as the RF distance between the true locus tree and true gene tree (column 2). *# leaves* is the number of leaves in a gene/locus tree. *# species* is the number of species labels in a gene/locus tree. *Mean # copies per species* is the number of time a species appears in a gene/locus tree averaged across all species. *# dups in LT* is the number of duplication events in a locus tree. # dups in GT is the number of tagged duplication vertices in a gene tree (using our definition of correct tagging). # dups Apro tag is the number of vertices in a gene tree tagged as duplications by Apro's rooting and tagging algorithm, for true and estimated gene trees (true/500bp/100bp). *GTEE* is the RF distance between the true and estimated gene tree (500bp/100bp). Note that the loss/duplication rate ratio is 0.

| Dup level | ILS level | # leaves | # species | Mean # copies per species | # dups in LT | # dups in GT | # dups in Apro tag true/500bp/100bp | GTEE 500bp/100bp |
|---|---|---|---|---|---|---|---|---|
| 0 | 0.00 | 26 | 26 | 1 | 0 | 0 | 0/NA/NA | 0.28/0.53 |
| | 0.21 | 26 | 26 | 1 | 0 | 0 | 0/NA/NA | 0.22/0.51 |
| | 0.50 | 26 | 26 | 1 | 0 | 0 | 0/NA/NA | 0.22/0.51 |
| | 0.71 | 26 | 26 | 1 | 0 | 0 | 0/NA/NA | 0.23/0.50 |
| 1 | 0.00 | 54.07 | 26 | 2.08 | 4.50 | 4.50 | 4.50/5.35/6.57 | 0.21/0.48 |
| | 0.19 | 48.76 | 26 | 1.88 | 3.77 | 3.99 | 3.91/4.61/5.55 | 0.18/0.47 |
| | 0.48 | 47.49 | 26 | 1.83 | 3.70 | 4.30 | 4.09/4.48/5.29 | 0.23/0.51 |
| | 0.68 | 48.80 | 26 | 1.88 | 3.77 | 4.79 | 4.42/5.43/6.39 | 0.20/0.48 |
| 2 | 0.00 | 67.92 | 26 | 2.61 | 7.39 | 7.39 | 7.39/8.28/10.20 | 0.19/0.44 |
| | 0.19 | 70.18 | 26 | 2.70 | 7.68 | 8.12 | 8.05/8.27/10.25 | 0.20/0.48 |
| | 0.46 | 64.21 | 26 | 2.47 | 6.92 | 8.03 | 7.80/8.19/9.69 | 0.19/0.48 |
| | 0.65 | 70.13 | 26 | 2.70 | 7.62 | 9.56 | 9.18/9.54/11.40 | 0.20/0.48 |
| 5 | 0.00 | 145.65 | 26 | 5.60 | 22.98 | 22.98 | 22.98/27.57/34.93 | 0.18/0.46 |
| | 0.19 | 151.45 | 26 | 5.83 | 24.21 | 25.80 | 25.74/25.65/31.90 | 0.18/0.46 |
| | 0.43 | 152.88 | 26 | 5.88 | 26.45 | 30.59 | 30.39/30.46/36.30 | 0.19/0.48 |
| | 0.62 | 150.59 | 26 | 5.79 | 24.72 | 30.59 | 30.25/28.49/33.03 | 0.20/0.49 |

Table S3: **D2 Summary Statistics.** All summary statistics are averaged across all 2000 gene trees and/or locus trees and then averaged across all 25 replicates. Columns are defined in Table S2, except *# loss in LT*, which is the number of loss events in a locus tree. Note that the population size was set to 1.9e+8, which corresponds ILS=0.5 for the model condition with no duplications or losses.

| Dup level | Loss/dup ratio | ILS level | # leaves | # species | Mean # copies per species | # dups in GT | # dups in LT | # loss in LT |
|---|---|---|---|---|---|---|---|---|
| 1 | 0.0 | 0.48 | 47.49 | 26 | 1.83 | 4.30 | 3.70 | 0.00 |
| | 0.5 | 0.45 | 39.59 | 22.45 | 1.78 | 3.63 | 3.46 | 1.59 |
| | 1.0 | 0.41 | 32.22 | 19.63 | 1.67 | 2.95 | 3.09 | 2.69 |
| 2 | 0.0 | 0.47 | 64.21 | 26 | 2.47 | 8.03 | 6.92 | 0.00 |
| | 0.5 | 0.42 | 49.20 | 21.10 | 2.35 | 6.13 | 6.11 | 2.83 |
| | 1.0 | 0.39 | 35.36 | 18.00 | 2.00 | 4.39 | 4.98 | 4.32 |
| 5 | 0.0 | 0.43 | 152.88 | 26 | 5.88 | 30.59 | 26.45 | 0.00 |
| | 0.5 | 0.39 | 69.76 | 19.44 | 3.59 | 12.91 | 13.37 | 6.36 |
| | 1.0 | 0.35 | 43.07 | 15.67 | 2.77 | 8.14 | 10.58 | 9.34 |

##### 1.3 Plant Data

The data used for the analysis on plant data can be downloaded at the https://github.com/chaoszhang/A-pro_data/blob/master/1kp/. We used two files:

- 1kp-c12-genetrees.tre : multi-copy gene trees
- astral-c12-renamed-induced-figtree.tre : ASTRAL species tree estimated from single-copy genes

### 2 Gene Tree Tagging

We gave TQMC-pro rooted and tagged gene trees as input, either the true ones or those computed by A-Pro v1.24.3.8 (download: https://github.com/chaoszhang/ASTER) with the following command:

```
astral-pro \
    -T \
    -i <input multilabeled tree> \
    -o <output tree> \
    -a <mapping file>
```

### 3 Species Tree Estimation

#### 3.1 Simulated Data Sets

The gene trees from SimPhy had leaves labeled accoding to the species, gene, and locus, so we created multi-labeled trees by keeping only the species name (i.e., we removed the part of the label following the first dash e.g. 0_0_0 became 0). We then estimated species trees from multi-labeled gene trees with the following software packages and commands.

**A-multi v1.24.4.8** (download: https://github.com/chaoszhang/ASTER) was run using the following command:

```
astral \
    -i <input multilabeled tree> \
    -o <output tree>
```

**TQMC v3.0.5 (commit: da4c498)** (download: https://github.com/molloy-lab/TREE-QMC) was run using the following command:

```
tree-qmc \
    -i <input multilabeled tree> \
    -o <output tree> \
    --norm_atax <Normalization scheme>
```

where normalization scheme was set to either 0 or 2.

**A-Pro v1.24.3.8** (download: https://github.com/chaoszhang/ASTER) was run using the following command:

```
astral-pro \
    -i <input multilabeled tree> \
    -o <output tree>
```

**TQMC-Pro commit: 194dab0** (download: https://github.com/molloy-lab/TREE-QMC/tree/tqmc-pro) was run using the following command:

```
tree-qmc \
    --gdl \
    --tagged \
    -i <input tagged multilabeled tree> \
    -o <output tree> \
    --norm_atax 0
```

#### 3.2 Plant Data Set

The gene trees for the plant data set were already multi-labeled by the species set. We estimated species trees from multi-labeled gene trees with the following software packages and commands.

**A-multi v1.24.4.8** was run using the following command:

```
astral -i <input tagged multilabeled tree> \
       --root "Uronema" \
       -u 1 \
       -o <output tree>
```

**A-Pro v1.24.3.8** was run using the following command:

```
astral-pro -i <input tagged multilabeled tree> \
           --root "Uronema" \
           -u 1 \
           -o <output tree>
```

**TQMC-Pro commit: 194dab0** was run using the following command:

```
tree-qmc \
    --gdl \
    --tagged \
    -i <input tagged multilabeled tree> \
    -o <output tree> \
    --norm_atax 0
```

We scored the TQMC-pro tree and the A-multi tree with A-pro using the following command:

```
astral-pro -C \
           -c <input species tree> \
           -i <input gene trees>
```

which also produced branch support annotations that we used for the TQMC-pro tree. Lastly, we scored the TQMC-pro tree and the A-pro tree with A-multi using the same command.

#### 4 Supplemental Results

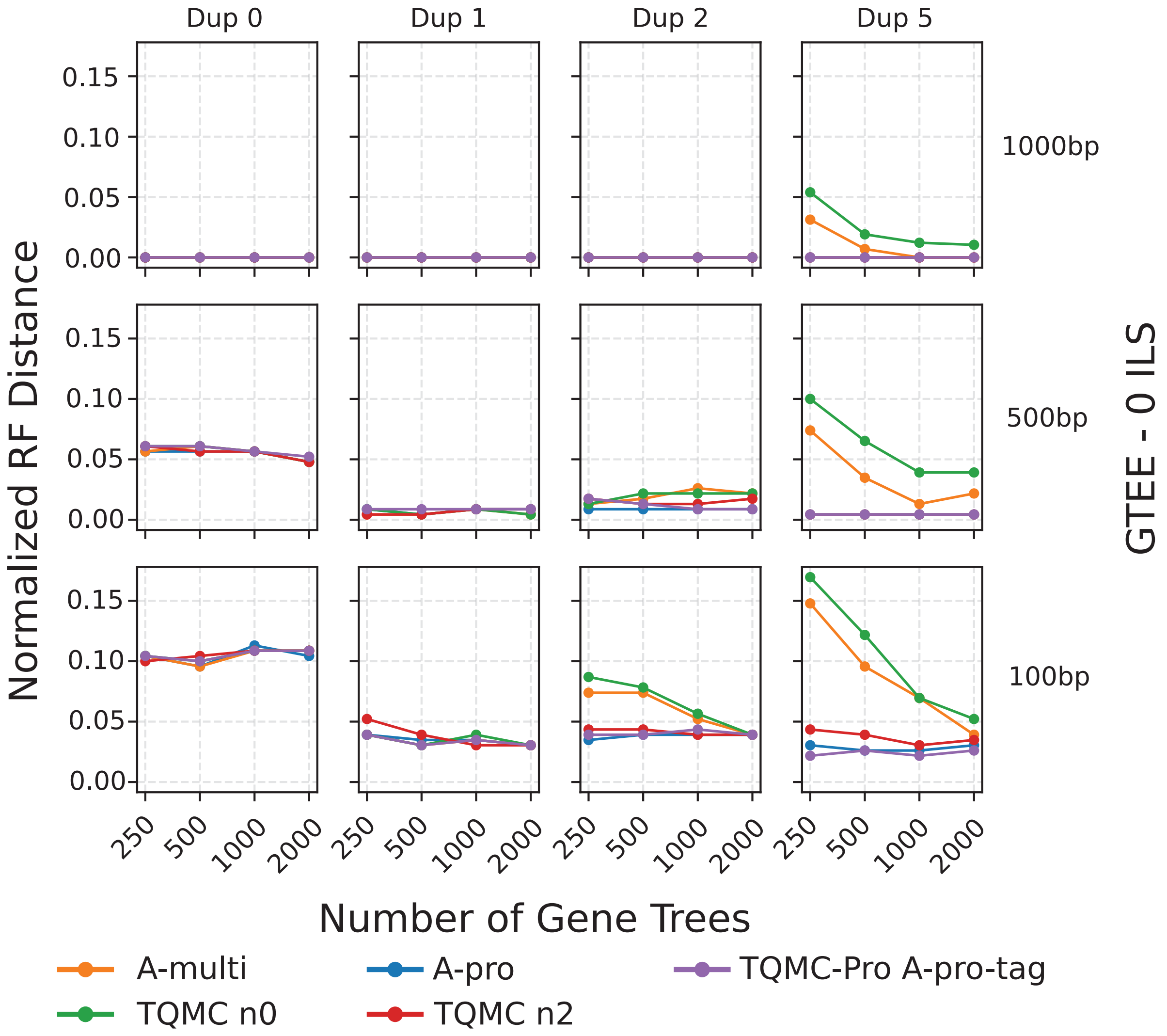

Figure S1: **Species tree error varying gene tree estimation error (ILS=0.0 and loss/dup ratio=0).** Dots are averages over 25 replicates for true gene trees and 10 replicates for estimated gene trees. Note that there is a strange trend for duplication level 0, where increasing the number of genes does not decrease species tree estimation error. Investigating this further, we found error was generally low across all except for a few outlier replicates; therefore, we conjecture that increasing the number of replicates from 10 to 25 (as was used for true gene trees) may have driven the error lower as in the duplication level 1 condition.

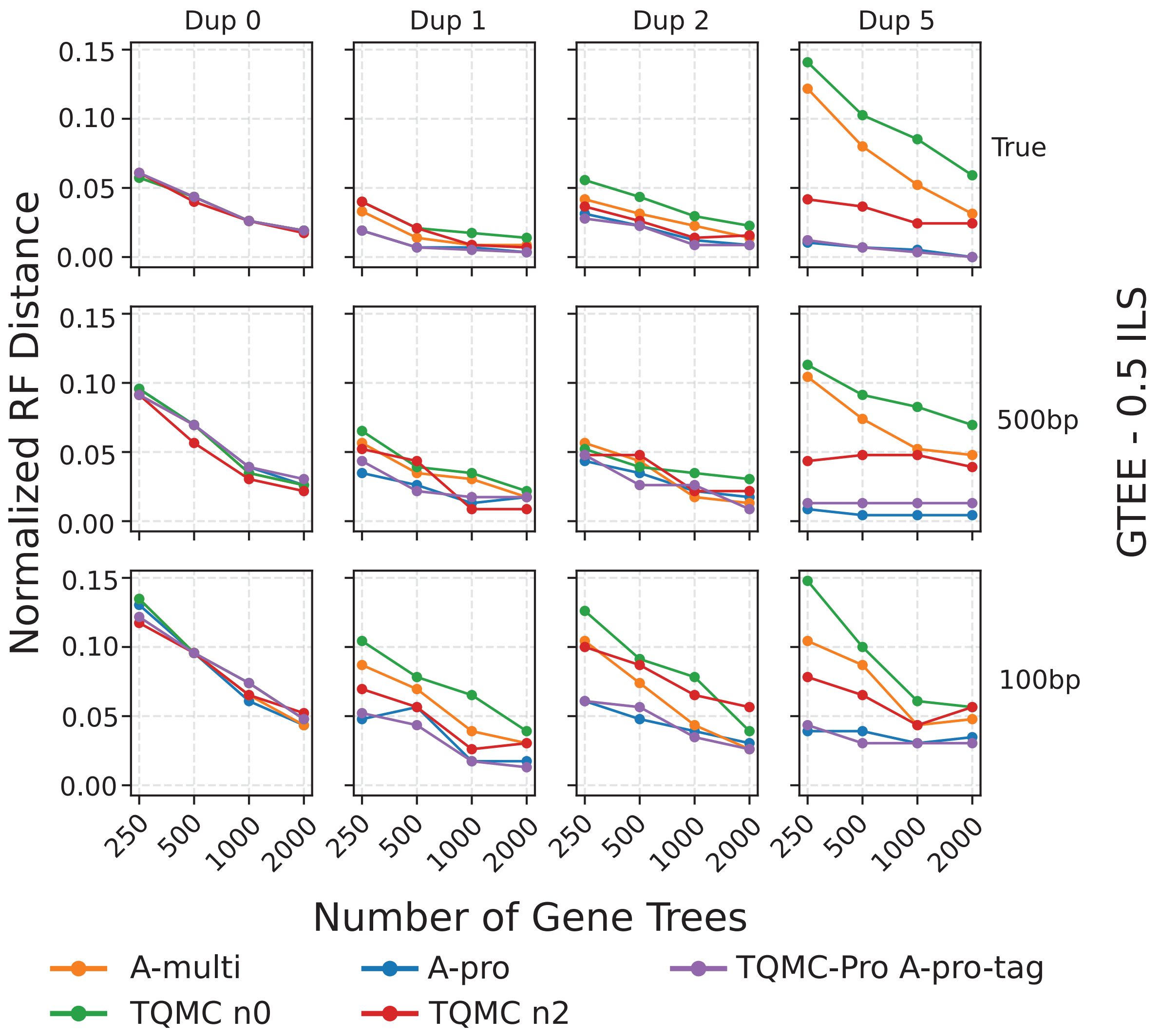

Figure S2: **Species tree error varying gene tree estimation error (ILS=0.5 and loss/dup ratio=0).**

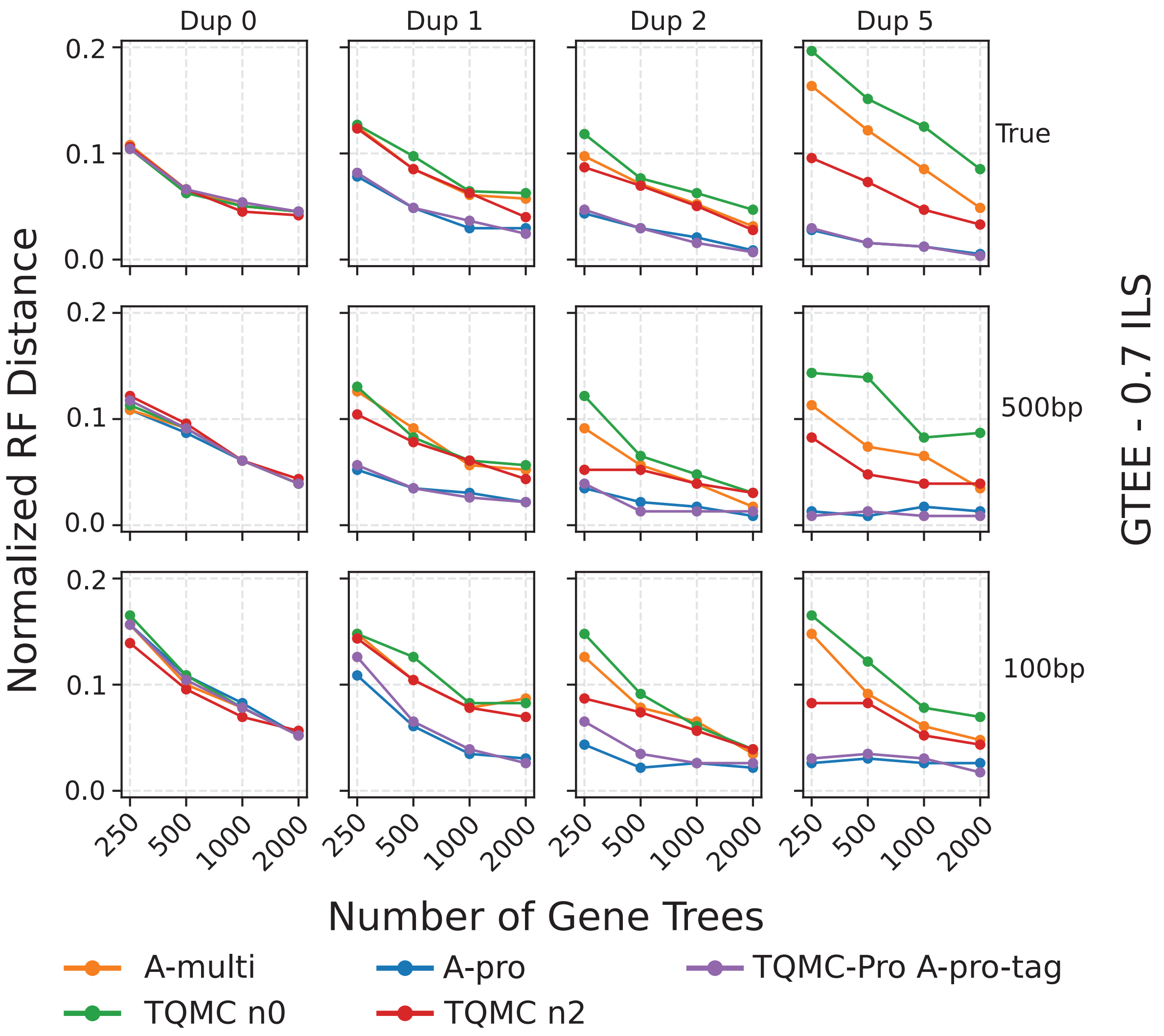

Figure S3: **Species tree error varying gene tree estimation error (ILS=0.7 and loss/dup ratio=0).**

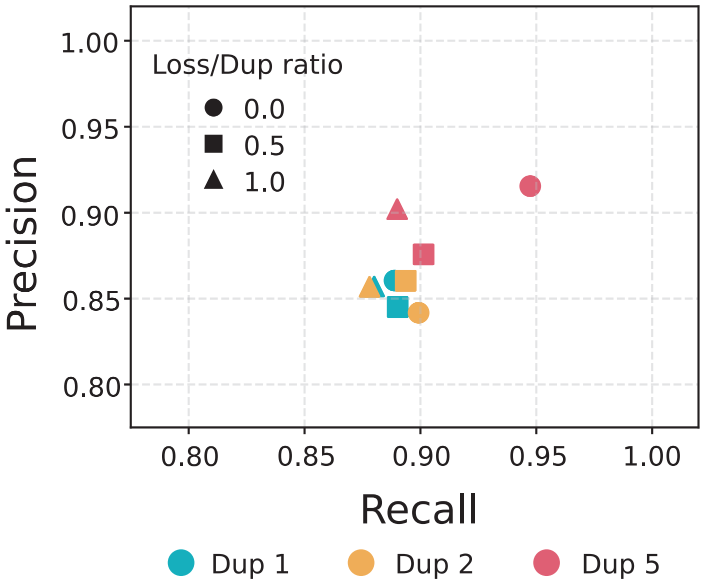

Figure S4: **Accuracy of Apro's gene tree tagging varying loss/duplication rate ratios (ILS=0.5).**

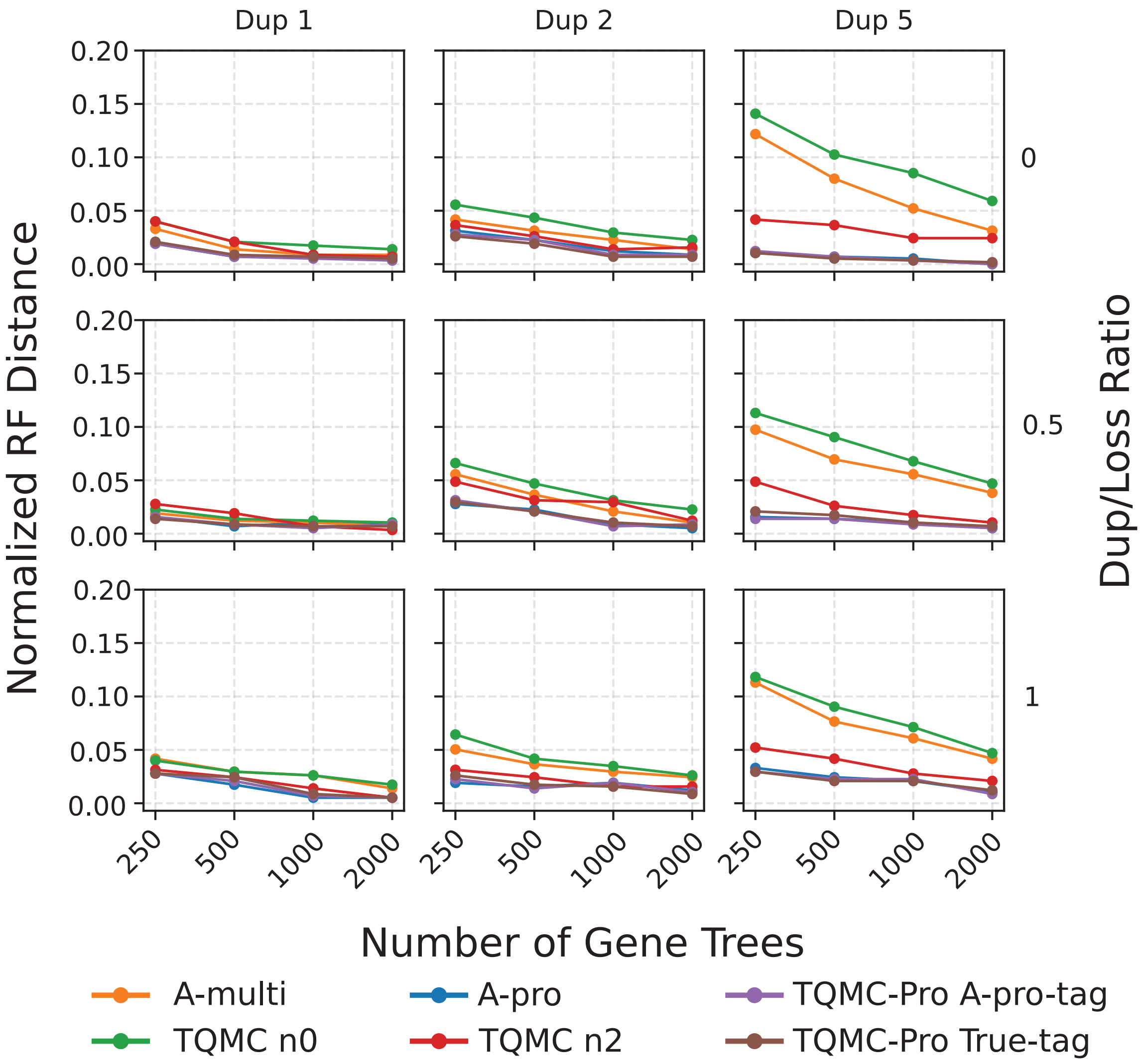

Figure S5: **Species tree error varying loss/duplication rate ratios (ILS=0.5).**

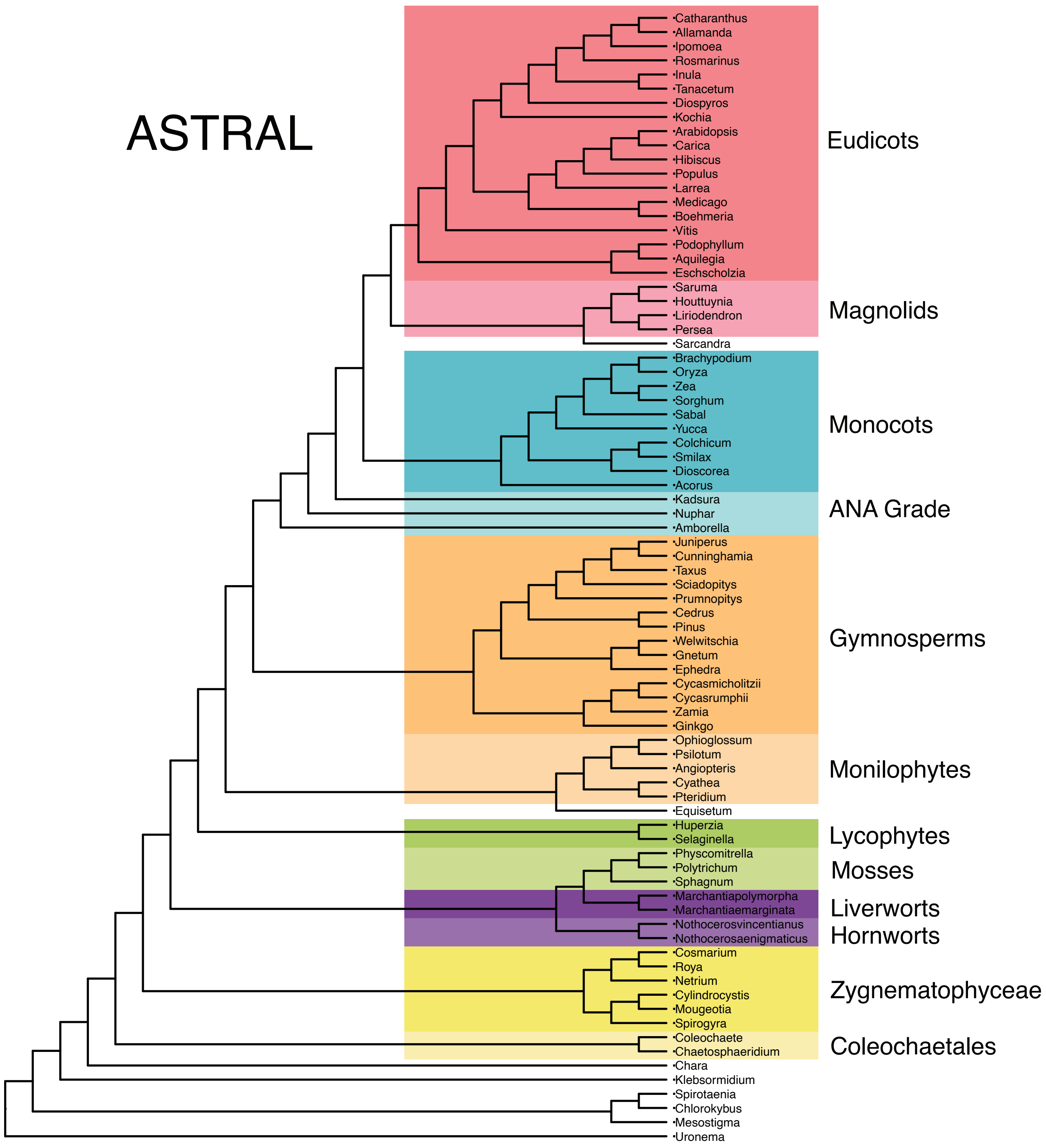

Figure S6: ASTRAL plant tree estimated from single-copy genes
